# Cross-regulation of (p)ppGpp and c-di-GMP pathways controls a cell-cycle transition

**DOI:** 10.1101/2025.08.22.671821

**Authors:** Corentin Jaboulay, Chloé Dugelay, José P.S. Maio, Katell Beillevaire, Philippe Vogeleer, Daniel Vinella, Fabien Létisse, Régis Hallez, Boyuan Wang, Laurent Terradot, Mathilde Guzzo

## Abstract

Bacteria must adapt their physiology to changing environments, a process often orchestrated by nucleotide messengers such as (p)ppGpp and cyclic di-GMP (c-di-GMP). While (p)ppGpp promotes survival under nutrient stress by suppressing growth, c-di-GMP supports sessile lifestyles and cell differentiation. In the α-proteobacterium *Caulobacter crescentus*, these messengers have opposite effects on the cell cycle, suggesting potential cross-regulations. Here, we uncover direct molecular cross-talk between (p)ppGpp and c-di-GMP signaling pathways. Using biochemical, structural and genetic approaches, we show that (p)ppGpp binds to the diguanylate cyclase PleD and inhibits c-di-GMP synthesis by competing with GTP at the active site. This interaction prevents c-di-GMP accumulation under nutrient-limiting conditions, linking metabolic stress to inhibition of developmental progression. Our findings reveal a new paradigm for second messenger interplay, in which one messenger directly suppresses synthesis of another. This work provides insight into how bacteria integrate metabolic and developmental cues through small-molecule cross-talk to guide adaptive behavior.

## Introduction

All living organisms constantly sense and respond to their environment to adapt to changing conditions. In bacteria, these responses can lead to a temporary arrest of growth, followed by renewed proliferation when conditions improve. These adaptive responses are orchestrated by complex signaling networks that rely on a wide array of regulatory molecules. Among them, nucleotide-based second messengers have emerged as central coordinators of bacterial physiology, integrating environmental cues to regulate a broad spectrum of cellular processes.

Molecular cross-talk between small signaling molecules is increasingly recognized as a critical mechanism for integrating multiple environmental cues. Several mechanisms have already been identified, including competition for common molecular targets, direct allosteric regulation of synthetic or degradative enzymes or metabolic competition (Fung et al., 2023). One example of small-molecule cross-talk is the binding of c-di-AMP to DarB/CbpB, which prevents DarB/CbpB from activating the *Bacillus subtilis* Rel synthetase/hydrolase (Ainelo et al., 2023; Krüger et al., 2021; Peterson et al., 2020), thereby inhibiting (p)ppGpp synthesis. Conversely, (p)ppGpp inhibits c-di-AMP degradation by targeting the phosphodiesterase GdpP (Corrigan et al., 2015; Rao et al., 2010), although this mechanism has not yet been characterized *in vitro*. In *Pseudomonas aeruginosa*, (p)ppGpp limits c-di-GMP synthesis by regulating purine salvage pathways (Kennelly et al., 2024).

Two of the most extensively studied nucleotide messengers are guanosine tetra- and pentaphosphate (collectively named (p)ppGpp) and cyclic di-GMP (c-di-GMP), each of which governs distinct, though sometimes overlapping, aspects of bacterial adaptation. (p)ppGpp is synthesized from GDP (ppGpp) or GTP (pppGpp) and ATP by members of the Rel/SpoT Homolog (RSH) family (Atkinson et al., 2011; Ronneau & Hallez, 2019)(Fig. 1A) in response to environmental stresses, particularly nutrient limitation (Battesti & Bouveret, 2006, 2009; Das et al., 2009; Potrykus & Cashel, 2008; Ronneau et al., 2016, 2019). Acting as a global regulator, (p)ppGpp represses growth-related processes and promotes survival, modulating transcription, translation, DNA replication, and metabolism (Steinchen et al., 2020). This conserved alarmone plays key roles in most bacterial species, as well as in plant chloroplasts and mitochondria (Atkinson et al., 2011; Ito et al., 2020). Conversely, c-di-GMP governs transitions between motile and sessile lifestyles. It is synthesized from GTP by diguanylate cyclases (DGCs) containing GGDEF domains (Fig. 1A) and degraded by phosphodiesterases (PDEs) with EAL or HD-GYP domains (Ryjenkov et al., 2005; Tal et al., 1998). High levels of c-di-GMP promote surface attachment, biofilm formation, and cellular differentiation, while low levels support motility and planktonic growth (Jenal et al., 2017).

**Figure 1:**
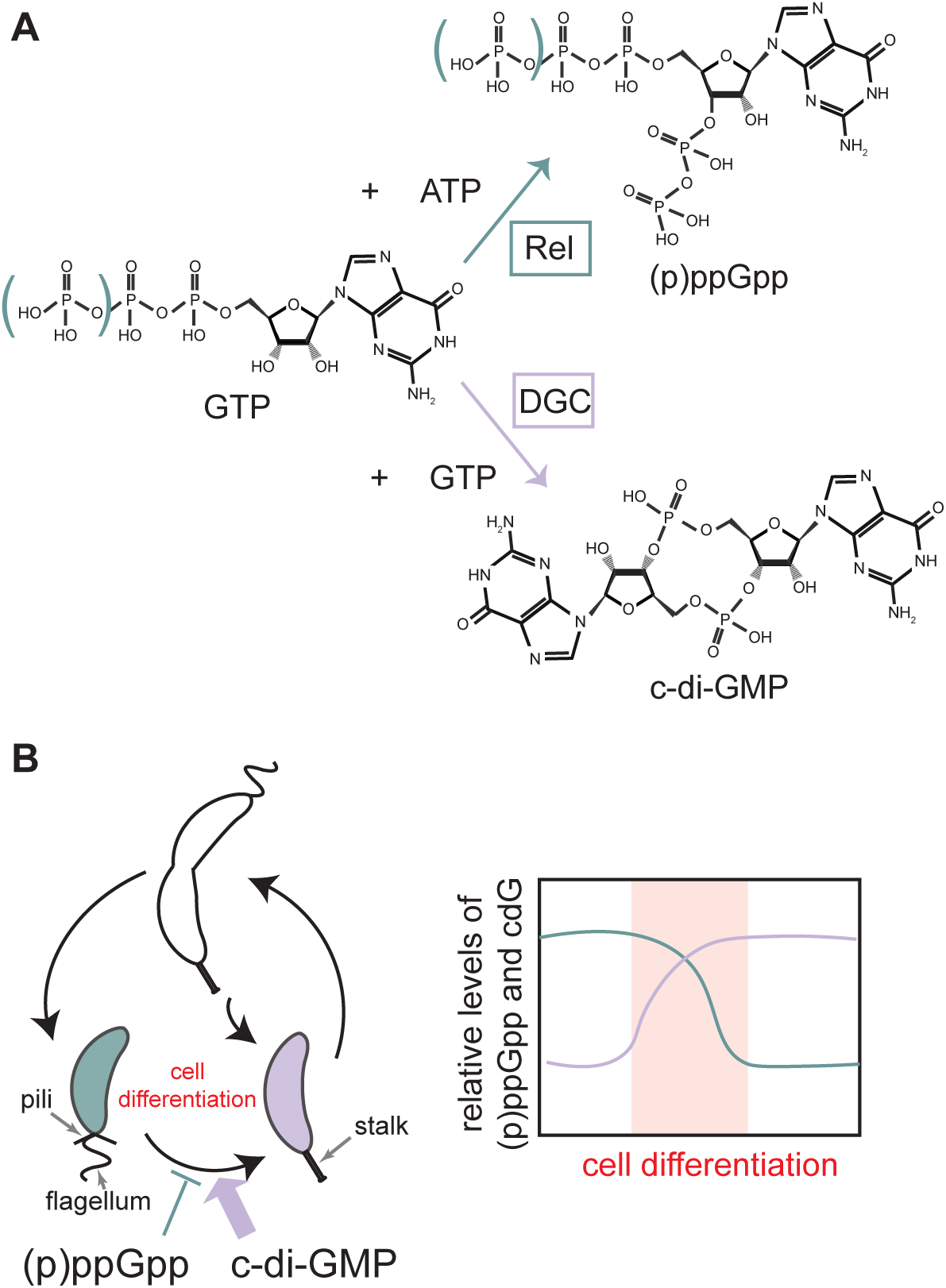
(p)ppGpp/c-di-GMP antagonism. (A) *Top:* Schematic of *Caulobacter crescentus* cell-cycle. Cells divide asymmetrically generating a swarmer daughter cell (with indicated flagellum and pili) and a stalked daughter cell (with indicated stalk). (p)ppGpp (green) and c-di-GMP (purple) accumulate at different steps of the cell-cycle and have antagonistic activities on cell differentiation respectively blocking and triggering transition from swarmer-to-stalked cell. *Bottom:* (p)ppGpp and c-di-GMP predicted oscillations during the cell-cycle. (B) Diagrams of enzymatic reactions leading to the synthesis of (p)ppGpp and c-di-GMP catalyzed by Rel enzyme or diguanylate cyclases (DGC) respectively. Both are made from guanosine-based nucleotides (GTP or GDP).

The α-proteobacterium *Caulobacter crescentus* provides a powerful model to investigate how such signalling pathways integrate environmental information to control developmental transitions. *C. crescentus* undergoes a well-characterized asymmetric cell division, producing a motile, non-replicative swarmer cell in G1 phase and a sessile, replicative stalked cell in S phase (Barrows & Goley, 2023; Collier, 2019)(Fig. 1B). Cell-cycle progression requires the swarmer cell to differentiate into a stalked cell. This transition is tightly controlled by (p)ppGpp and c-di-GMP: (p)ppGpp accumulates under carbon starvation, carbon exhaustion, nitrogen starvation or glutamine deprivation (Boutte et al., 2012; Boutte & Crosson, 2011; Britos et al., 2011; Hallgren et al., 2023; Lesley & Shapiro, 2008; Ronneau et al., 2016) and prolongs the G1 phase (Ronneau et al., 2016), blocking differentiation. Conversely, under favorable conditions, rising levels of c-di-GMP promote the swarmer-to-stalked transition and S-phase entry (Abel et al., 2013; Duerig et al., 2009; Lori et al., 2015) mainly through the activation of the PleD diguanylate cyclase (Christen et al., 2010; Kaczmarczyk et al., 2020, 2024; Paul et al., 2007, 2008), reinforcing their opposing roles in coupling environmental sensing to proliferation (Fig. 1B). Both second messengers are derived from GTP (or GDP for ppGpp) (Fig. 1A), suggesting a need for tight metabolic prioritization and temporal separation of their synthesis during the cell cycle.

Due to its genetic tractability and well-characterized cell cycle, *C. crescentus* offers a powerful model for dissecting nucleotide signaling cross-talk. In this study, we uncover a direct molecular antagonism between (p)ppGpp and c-di-GMP. Using biochemical, structural, and genetic approaches, we show that (p)ppGpp binds directly to the active site of the diguanylate cyclase PleD and competes with its GTP substrate, thereby inhibiting c-di-GMP synthesis. This mechanism reveals a direct link between nutrient stress signaling and developmental regulation in *C. crescentus* and provides new insight into how bacteria integrate multiple inputs to fine-tune growth and differentiation.

## Results

### (p)ppGpp directly binds to a c-di-GMP-producing enzyme

To investigate potential molecular connections between (p)ppGpp and c-di-GMP signaling, we employed a ppGpp capture-compound strategy(Wang et al., 2019) to search for ppGpp-binding proteins in *C. crescentus* (Fig. 2AB, Fig. S1). This approach uses a trifunctional molecule where ppGpp is derivatized to include a cross-linkable moiety and a biotin handle(Wang et al., 2019). The cross-linkable ppGpp variants were incubated with two *C. crescentus* cell lysates: one grown in minimal medium with ^15^N-labeled ammonium chloride and another in standard minimal medium (see Methods, Fig. 2A). Unlabeled lysates were outcompeted with excess of unmodified ppGpp. Labeling with ^15^N allows the calculation of heavy-to-light peptide ratios, thereby identifying proteins that remain bound to ppGpp even in the presence of excess unmodified competitor. Following cross-linking, proteins were isolated via streptavidin pull-down, digested with trypsin, and analyzed by mass spectrometry (see Methods, Fig 2AB). This screen identified homologs of known ppGpp targets in other bacteria, including hypoxanthine-guanine phosphoribosyltransferase (HPRT), a MutT-like protein, and the HflX GTPase (Bennison et al., 2021; Corrigan et al., 2016; Wang et al., 2019; Yang et al., n.d.; Zhang et al., 2018)(Fig. 2C). Additionally, we identified PleD, a diguanylate cyclase (DGC), as a potential ppGpp-binding protein with an average heavy-to-light ratio of 11 (log₂ H/L = 3.5, Fig. 2C, see Methods).

**Figure 2:**
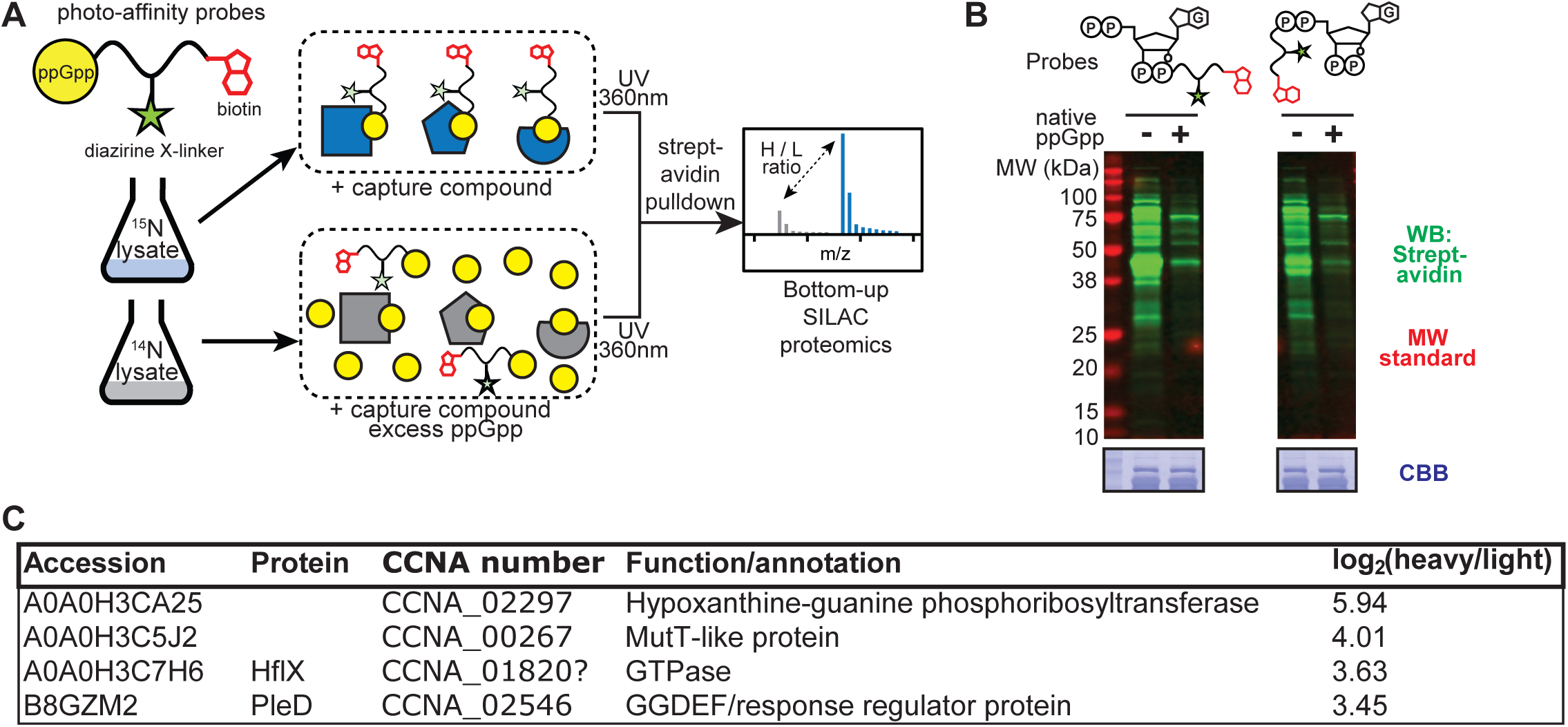
ppGpp-target identification in *Caulobacter crescentus*. (A) Chemical proteomic approach for ppGpp-target identification based on metabolic universal ^15^N-labeling of *C. crescentus* in M2G medium. (B) *C. crescentus* proteome tagged by indicated probes visualized using streptavidin western blotting. See Figure S1 for probe structure and synthesis. Lysate containing 100 μM probe (ppGpp) or 100 μM probe and 5 mM native ppGpp (+ ppGpp) was illuminated with 360-nM UV. Aliquots containing 10 μg protein were resolved on 12% SDS-PAGE for immunoblotting (C) Putative ppGpp-binding proteins identified. Shown are homologs of known ppGpp targets and the diguanylate cyclase PleD investigated in this study.

PleD is a response regulator that consists of a phosphorylatable N-terminal receiver domain (D1), a receiver-like adaptor domain (D2) and a DGC domain. PleD catalyzes the synthesis of c-di-GMP from two GTP molecules in its active site within the DGC domain. Phosphorylation of PleD at aspartate D53 of the D1 domain enhances PleD activity, likely by promoting dimerization(Wassmann et al., 2007), and is mediated by the PleC and DivJ histidine kinases during *C. crescentus* differentiation(Chong et al., 2024; Kaczmarczyk et al., 2020; Paul et al., 2004). PleD activation during cell differentiation triggers the accumulation of intracellular c-di-GMP which drives the morphological changes from swarmer to stalked cell and the transition to a replicative cell. Since accumulation of (p)ppGpp has the opposite effect of blocking cell differentiation in *C. crescentus,* a potential PleD-(p)ppGpp interaction suggests an antagonistic cross-talk between c-di-GMP and (p)ppGpp signaling networks, that we explore below.

### (p)ppGpp directly binds PleD and inhibits its diguanylate cyclase activity

To investigate if PleD is a (p)ppGpp target, we purified recombinant *C. crescentus* PleD from *E. coli* (Chan et al., 2004) and assessed binding to ppGpp and pppGpp using nano differential scanning fluorimetry (nanoDSF) and isothermal titration calorimetry (ITC) (Table S1-S2). Of note, during purification, a fraction of DGCs was recovered with c-di-GMP bound, most likely at their inhibitory sites (I-site) (Fig. S2A). Similar observations have been reported for purified GGDEF enzymes (Chan et al., 2004; De et al., 2009; Wassmann et al., 2007), although the extent varies. NanoDSF assays revealed significant increases in the melting temperature of PleD in the presence of both ppGpp and pppGpp (Fig. S2BC). Notably, pppGpp induced a larger shift (ΔTm = 11.52°C ± 0.23) than ppGpp (ΔTm = 6.332°C ± 2.323) (Fig. S2BC, Table S2). No significant shift was observed with another purine nucleotide, ATP, highlighting nucleotide specificity (Fig. S2BC). ITC revealed a 1:1 binding stoichiometry for both ppGpp (K_D_ = 45.4 µM ± 22.3) and pppGpp (K_D_ = 9.31 µM ± 1.16) (Fig. 3A). Interestingly, pppGpp bound PleD with an affinity comparable to GMPCPP, a non-hydrolyzable GTP analog and substrate mimic (Fig. 3A, Fig. S3A). These data reveal a strong and specific interaction between PleD and (p)ppGpp, with tighter binding by pppGpp than ppGpp.

**Figure 3:**
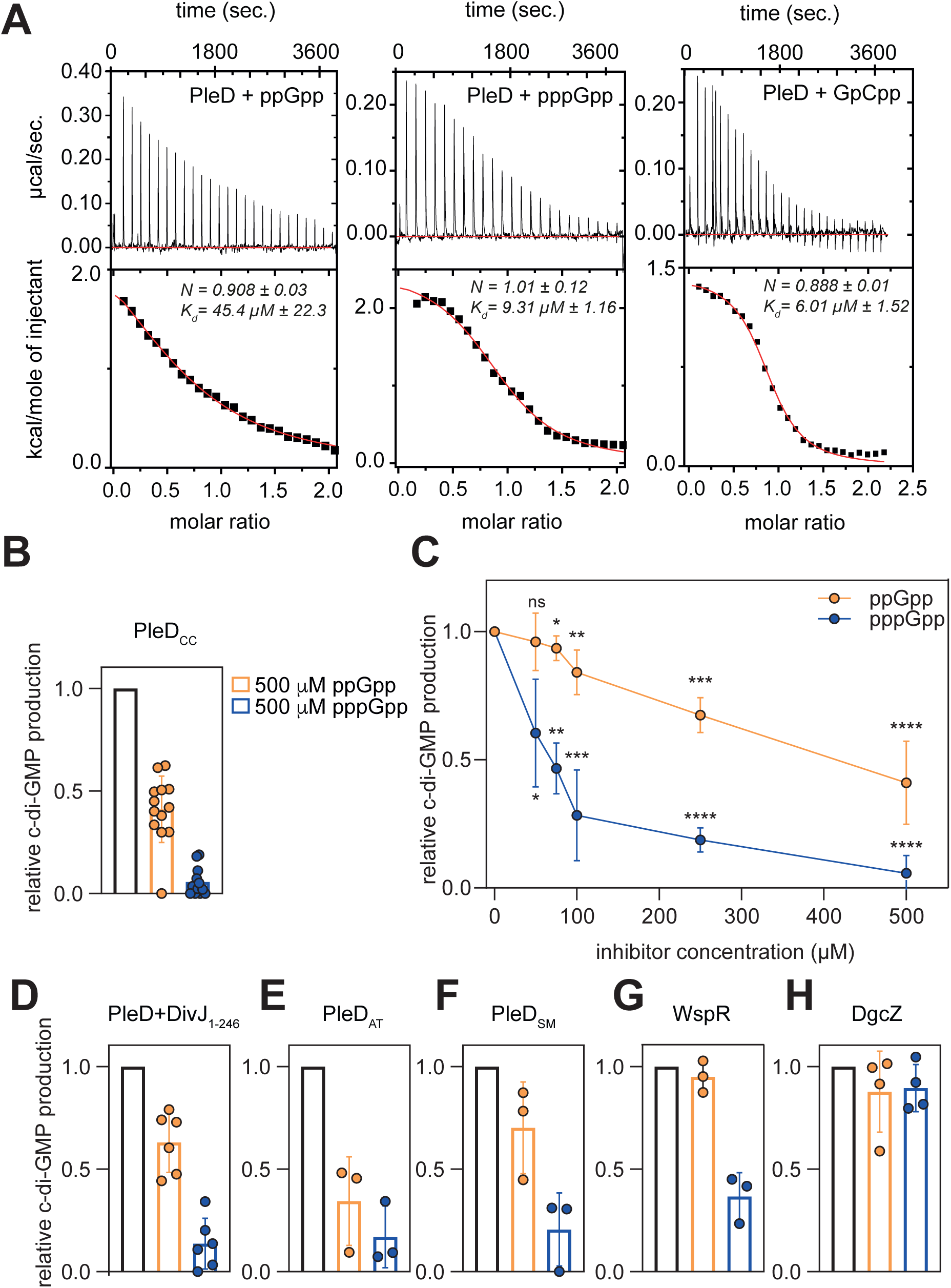
Inhibition of PleD’s activity by ppGpp and pppGpp through direct binding. (A) ITC traces (top) and fitted isotherms (bottom) for the titration of 100 µM PleD with 1 mM ppGpp, pppGpp, or GpCpp (left to right). K_D_: dissociation constant; *N*: stoichiometry of the PleD-ligand complex. (B) *In vitro* enzymatic assay measuring c-di-GMP synthesis by PleD from *Caulobacter crescentus*. Reactions contain 40 µM PleD and 100 µM GTP, either alone (black) or with 500 µM ppGpp (orange) or pppGpp (blue). The relative c-di-GMP production shows the amount of c-di-GMP generated under each condition, normalized to the production observed in the absence of competitor. Data are expressed as mean ± SD. (C) Dose-response curves for c-di-GMP production, normalized to the control, with 40 µM PleD and 100 µM GTP in the presence of increasing concentrations of ppGpp (orange) or pppGpp (blue): 50 µM, 75 µM, 100 µM, 250 µM, or 500 µM. The relative c-di-GMP production was calculated by comparing each condition to the control where no competitor was present. Data represent mean ± SD from 3 independent experiments (for 50 µM, 75 µM, 100 µM and 250 µM conditions) and 12 independent experiments (for 500 µM condition). Asterisks indicate statistical significance (*= p <0.05; **= p <0.01; ***= p <0.001; ****= p <0.0001) based on one-sample Student’s t-test. (D) Same as (B), but for phospho-activated PleD pre-incubated with 10 µM DivJ_1-246_ and 200 µM ATP for 25 minutes prior to the addition of 500 µM ppGpp (orange) or pppGpp (blue). (E) Same as (B) but for PleD ortholog from *Agrobacterium tumefaciens* (PleD_AT_). (F) Same as (B) but for PleD ortholog from *Sinorhizobium meliloti* (PleD_SM_). (G) Same as (B) but for WspR, a PleD ortholog from *Pseudomonas aeruginosa*. (H) Same as (B) but for DgcZ from *Escherichia coli*.

To explore a potential role of (p)ppGpp on PleD’s enzymatic activity, we assessed PleD’s *in vitro* c-di-GMP production using reverse phase (or C18)-HPLC(Paul et al., 2004). In the presence of GTP at 100 µM, PleD produced c-di-GMP (Paul et al., 2004) (Fig. S3B). Addition of either ppGpp or pppGpp significantly impaired the enzymatic activity of PleD *in vitro* (Fig. 3BC). At a 1:5 GTP-to-ppGpp ratio, the c-di-GMP synthesis activity of PleD was reduced by more than 50% (Fig. 3BC). In the same conditions, pppGpp exhibited an even stronger inhibitory effect, nearly abolishing c-di-GMP production with over 90% inhibition (Fig. 3BC). Under varying alarmone concentrations in the reaction mixture, pppGpp demonstrated potent inhibition at a 1:1 ratio with GTP (Fig. 3C), and showed a greater efficacy in suppressing PleD activity compared to ppGpp at a similar ratio. These results indicate that pppGpp is a more effective inhibitor of PleD than ppGpp.

In contrast to (p)ppGpp, ATP had no detectable effect on PleD activity (Fig. S3A), underscoring the enzyme’s specificity for guanosine-derived nucleotides. Phosphorylation of PleD at residue D53, mediated by the truncated DivJ kinase *in vitro*(Paul et al., 2004), led to an approximately 2.5-fold increase in enzymatic activity (Fig. S3A), aligning with previous findings(Paul et al., 2004). Despite this activation, (p)ppGpp remained potent inhibitors (at a 1:5 GTP:(p)ppGpp ratio), significantly reducing the activity of PleD in its phosphorylated state (Fig. 3D, Fig. S3C). These findings emphasize that the inhibitory effect of (p)ppGpp operates independently of PleD’s phosphorylation status, indicating a robust and direct mechanism of action.

### Structural basis of PleD inhibition by (p)ppGpp

To gain insights into the mechanism of PleD’s inhibition by these alarmones, we solved PleD’s crystal structure in complex with ppGpp (PleD^ppGpp^) or pppGpp (PleD^pppGpp^) at resolutions of 2.9 Å and 3 Å, respectively (Table S3). The complexes crystalized in the space group P6_1_22, with four molecules per asymmetric unit, arranged as dimers of dimers (Fig. S4AB). In both structures, the overall conformation of the PleD dimer resembles the inactive conformation described in the PleD:c-di-GMP binary complex structure previously published (Chan et al., 2004) (Fig. 4A, Fig.S4C), albeit domain swapping between the D1-D2 and the DGC domain was observed in our structures. Compared to the inactive conformation, the stem α5 helices undergo a small conformational change with helices extended by 4-5 residues, reminiscent of what was observed upon activation by phosphorylation (Wassmann et al., 2007) (Fig. S4C). Of note, both the PleD^ppGpp^ and the PleD^pppGpp^ structures were solved in the presence of c-di-GMP, thus each monomer of the (p)ppGpp-PleD structures is bound by two cyclic di-GMP molecules at its I-site, *i.e.* between the DGC and the D2 domains(Chan et al., 2004) (Fig. 4AB). Each PleD DGC domain is bound by a (p)ppGpp and a Mg^2+^ ion in the active site (Fig. 4AB).

**Figure 4:**
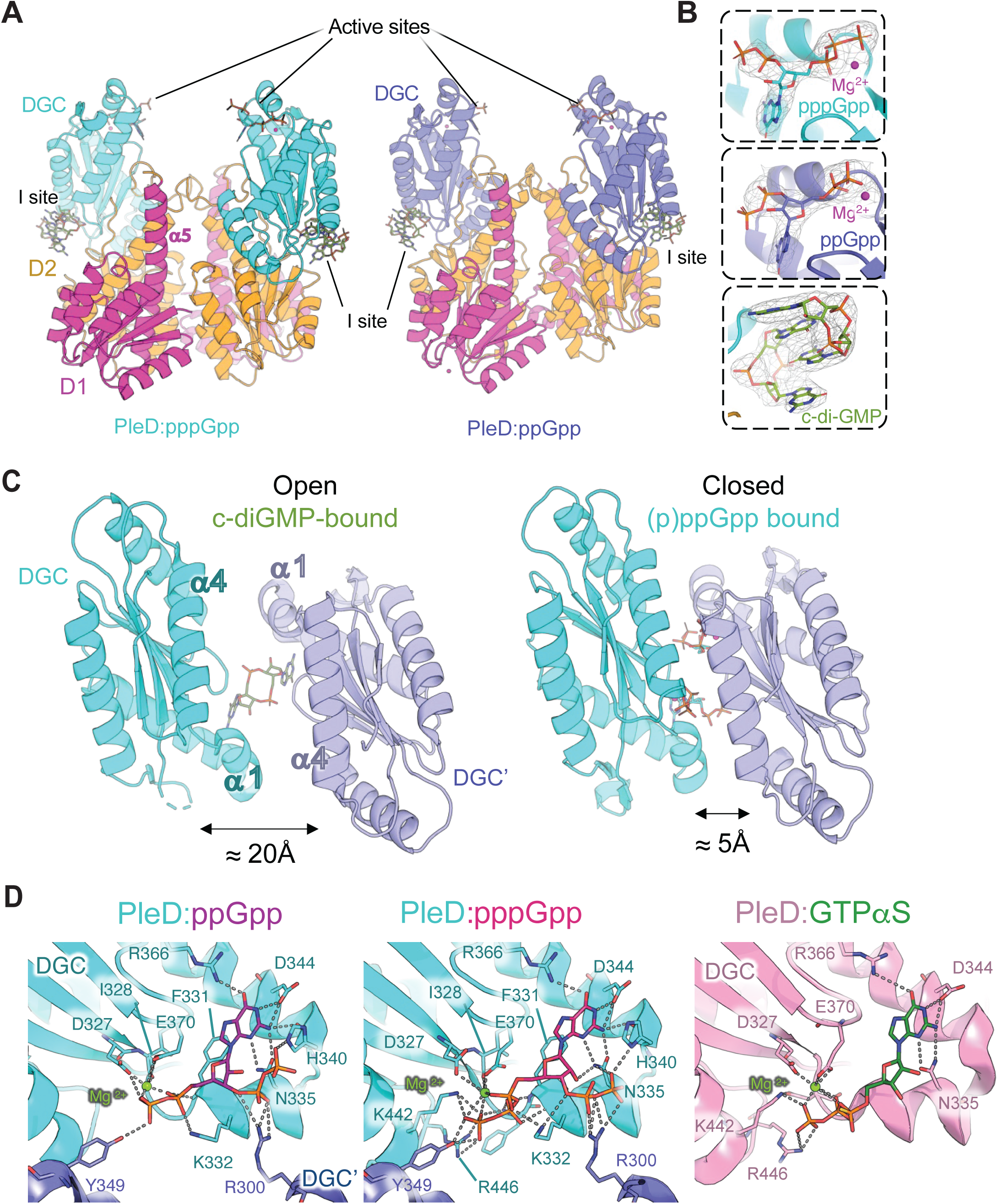
Structural insights into PleD inhibition by (p)ppGpp. (A) Ribbon representation of the crystal structure of PleD in complex with pppGpp (left) and ppGpp (right). The receiver domains D1 and D2 are colored in magenta an orange, respectively. The DGC domains are colored in cyan for PleD:pppGpp and slate for for PleD:ppGpp. The alarmones and c-di-GMP are represented as sticks. Carbones of alarmones are colored according to their DGC domain, c-di-GMP carbons are colored in green, oxygens are colored in red, phosphorus in orange and magnesium in magenta. The active and inhibitory sites (I-site) are indicated. (B) Representative close-up view of the alarmone and cyclic-di-GMP molecules colored as in A) with corresponding 2Fo-Fc electron density maps contoured at 1.5 sigma. (C) Comparison of the DGC domain’s position of cyclic-di GMP-bound PleD (PDB code 1W25, left) with the DGC domain’s position of pppGpp-bound PleD. (D) Comparison of the active site from PleD:GTPαS structural data (PDB code 2V0N) and the PleD:(p)ppGpp active sites. Ligands are shown in stick representation with carbons colored in purple (ppGpp), magenta (pppGpp) or green (GTPαS). The DGC domains contributing to the binding are colored in cyan and slate (alarmones) or pink (GTPαS). Magnesium atom is represented as a green sphere. Side chains involved in the binding are shown as sticks and hydrogen bonds are indicated by grey dashes.

The binding of the alarmones is at the interface of the dimer of dimers, bringing together two DGC domains, with each of them interacting with two (p)ppGpp (Fig. 4C, Fig. S4A-C). Consequently, these two DGC domains are in a “closed” conformation, contrasting with the open conformation identified in the inactive, product-bound PleD (Fig. 4C). This is illustrated by the difference of position of helices α4 (following the nomenclature in(Chan et al., 2004)) from the two DGC domains. While these are 20 Å away from each other in the product-bound form, they are less than 5 Å in the (p)ppGpp-bound structures (Fig. 4C). The binding mode of GDP and GTP moieties of ppGpp and pppGpp into the individual DGC domain is similar to the one described for the non-hydrolyzable GTP-αS(Wassmann et al., 2007) (Fig. 4D). The guanine moiety sits in a pocket formed by DGC α1 and α2 and is hydrogen-bonded by D344 and N335. F331 main chain and K332 side chain interact with the β-pyrophosphate (Fig. 4D) while I328 main chain, D327 and E370 side chains bind the magnesium ion (Fig. 4D). The 3’ pyrophosphates of both alarmones point away from the active site although density for the second phosphate was rather poor (Fig. 4AB). The α−phosphate interacts with N335 but also with R300 from the other DGC domain (Fig. 4D). As a consequence, the 3’ pyrophosphate might interact with H340 side chain but this interaction is not absolutely conserved in the four subunits of the asymmetric unit. This might be explained by the charge of the surface at this site being predominantly negative and thus unfavourable to binding of negatively charged pyrophosphate. Residue N335 has been shown to be important for PleD DGC activity(Chan et al., 2004; Christen et al., 2006). We also observed that residue R300, which is highly conserved in DGC domains (Fig. S5), plays an important role in PleD’s activity. Indeed, substituting R300 with either alanine or lysine significantly reduced enzymatic activity (Fig. S4D). Thus, (p)ppGpp binding reveals key residues involved in PleD’s catalytic function, potentially stabilizing intermediate reaction conformations that were not captured in previous PleD structures. The main difference between the binding of ppGpp and pppGpp occurs with pppGpp 5’ γ-phosphate being bound by R446 and also by Y439, two highly conserved residues in DGC domains and the latter from the second DGC α4 helix (Fig. 4CD, Fig. S5). This binding adds two hydrogen bonds, explaining the increased affinity of pppGpp for PleD and its higher inhibitory effect compared to ppGpp.

### Conservation of (p)ppGpp-mediated inhibition across PleD orthologs

Diguanylate cyclases are widely conserved across bacteria and in some eukaryota (Kawabe et al., 2023; Römling et al., 2013). The active site of the DGC domain is particularly conserved among DGC enzymes of different families (Fig. S5), with numerous conserved residues that have been shown to be involved in the enzymatic reaction and/or dimerization. As we showed that (p)ppGpp ligands can bind in the active site of the PleD_CC_ DGC, interact with conserved residues and affect its activity, we investigated the conservation of (p)ppGpp sensitivity in this enzyme family and beyond. We purified PleD orthologs from *Agrobacterium tumefaciens* and *Sinorhizobium meliloti*, both part of the α-proteobacteria class, and assessed their activities *in vitro*. In both cases, ppGpp and pppGpp strongly inhibited c-di-GMP production, with pppGpp showing greater potency (Fig. 3EF, Fig. S3DE). NanoDSF assays further confirmed PleD_AT_ binding to (p)ppGpp, indicating conserved interaction mechanisms (Fig. S2DE; Table S2).

We expanded these studies to WspR, a PleD ortholog from *Pseudomonas aeruginosa*, representing γ-proteobacteria. We observed that ppGpp had minimal impact on WspR activity while pppGpp inhibited more than 50% of its enzymatic function (Fig. 3G, Fig. S3F). In contrast, DgcZ, a diguanylate cyclase with a different domain organization showed no reduction in activity (Fig. 3H, Fig. S3G). These results suggest that (p)ppGpp-mediated inhibition is a conserved feature of PleD orthologs, with pppGpp exhibiting greater inhibitory efficacy and potential conserved activity beyond α-proteobacteria.

### (p)ppGpp inhibition of PleD activity prevents cell differentiation under glucose exhaustion in *C. crescentus*

Having established biochemically that (p)ppGpp directly inhibits PleD activity, we next examined the physiological relevance of this interaction *in vivo*. Under glucose exhaustion conditions (M2G_1/10_*), *C. crescentus* cells remain arrested at the swarmer, non-replicative stage of the cell cycle, preventing differentiation into stalked cells and the initiation of DNA replication until conditions become favorable again (Boutte et al., 2012)(Fig. S6A). Previous studies have shown that glucose exhaustion triggers the accumulation of ppGpp and pppGpp in *C. crescentus* cells (Boutte et al., 2012; Hallgren et al., 2023; Leslie et al., 2015). Based on our *in vitro* data, we propose that the starvation-induced accumulation of (p)ppGpp inhibits PleD-mediated c-di-GMP synthesis, thereby involved in blocking the swarmer-to-stalked cell transition.

To investigate this model *in vivo*, we quantified for the first time in *C. crescentus* the intracellular concentrations of ppGpp, pppGpp, c-di-GMP, GTP, and GDP in mixed populations of cells grown under nutrient-rich conditions (M2G), nitrogen starvation (P2G), carbon exhaustion (M2G_1/10_*), and carbon starvation (M2) using LC-MS (Fig. 5A–C, Fig. S6BC; Table S4; see Methods). Under nitrogen starvation, ppGpp levels increased to ∼85 µM, and to ∼55 µM under carbon starvation, compared to ∼30 µM in nutrient-replete conditions (Fig. 5A, Table S4). Interestingly, ppGpp levels remained largely unchanged under carbon exhaustion. By contrast, pppGpp concentrations rose markedly to ∼10 µM under carbon exhaustion, whereas they remained around ∼4 µM in all other tested conditions (Fig. 5B, Table S4). Notably, the intracellular concentrations of ppGpp and pppGpp are comparable to the K_D_ values determined by ITC (Fig. 3A), suggesting that PleD binding by (p)ppGpp may be finely tuned under physiological conditions.

**Figure 5:**
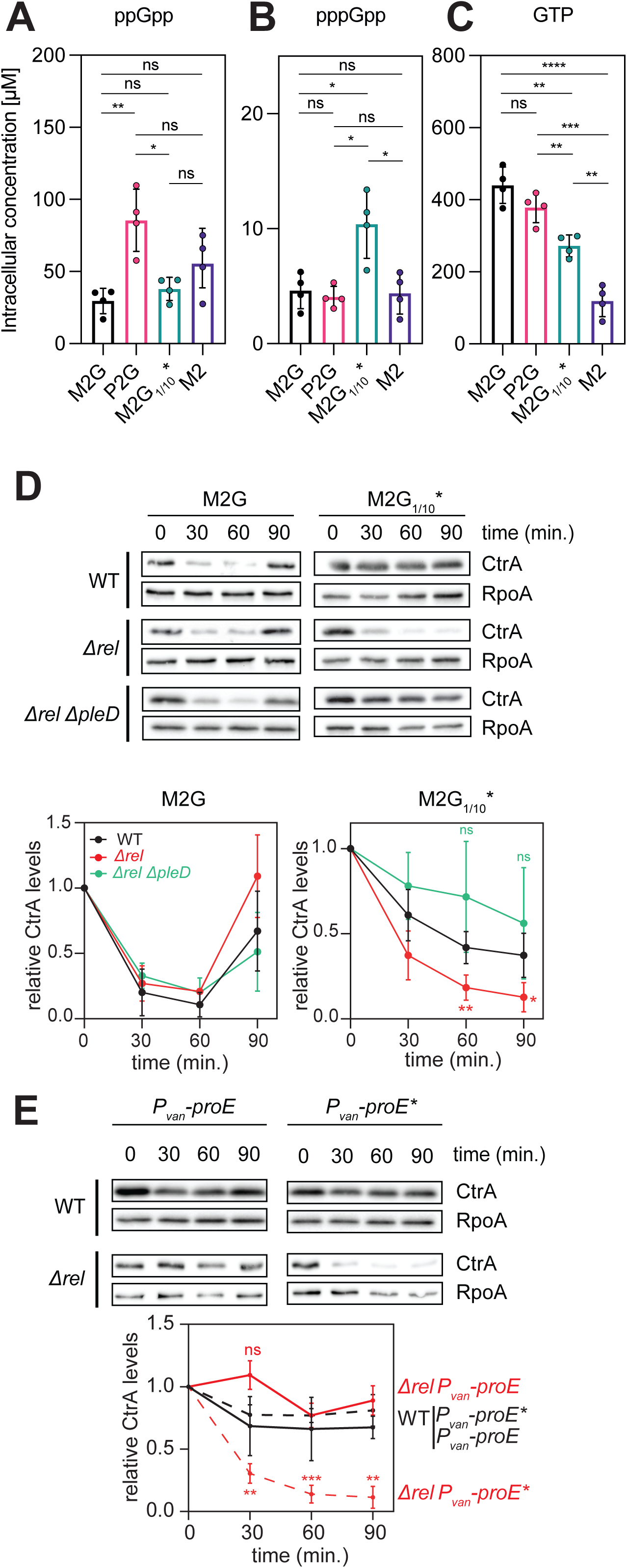
(p)ppGpp inhibition of PleD activity prevents cell differentiation under glucose starvation in *Caulobacter*. (A) Targeted metabolomics measuring the intracellular concentration of ppGpp in a asynchronous population of wild-type cells grown in M2G medium (black), P2G medium (pink), M2 with 0.02% glucose (M2G_1/10_*) (green) or M2 without glucose (purple). Data represent the mean ± SD of three independent biological replicates. Asterisks indicate statistical significance (** = p < 0.01; *** = p < 0.001) based on a Student’s t-test. (B) Same as (A) but representing pppGpp intracellular concentration. (C) Same as (A) but representing GTP intracellular concentration. (D) *Top:* Immunoblots showing CtrA protein levels at the indicated times post-synchronization in wild-type (WT), *rel* mutant, and *rel pleD* mutant strains grown in M2G medium or grown in M2 with 0.02% glucose spent medium (M2G_1/10_*). *Bottom:* Quantification of CtrA band intensity normalized to the RpoA loading control. Data represent the mean ± SD of three independent biological replicates. (E) Immunoblots showing CtrA protein levels at the indicated times post-synchronization in WT and *rel* mutant strains grown in M2 with 0.02% glucose spent medium (M2G_1/10_*), with ectopic expression of either the phosphodiesterase *proE* or its catalytically inactive mutant *proE_E328A_* (*proE\**) (induced with 100 µM vanillate post-synchronization) (left). Quantification of CtrA band intensity normalized to the RpoA loading control (right). Data represent the mean ± SD of three independent biological replicates. Asterisks indicate statistical significance (** = p < 0.01; *** = p < 0.001) based on a Student’s t-test.

GTP concentrations decreased from ∼440 µM in nutrient-rich conditions to ∼271 µM under carbon starvation and ∼119 µM under carbon exhaustion (Fig. 5C, Table S4), consistent with a global metabolic downshift under nutrient depletion. GDP levels also decreased similarly to the GTP concentrations (Fig. S6B, Table S4). In contrast, c-di-GMP levels remained largely unchanged across all tested conditions (Fig. S6C, Table S4), despite marked alterations in GTP and (p)ppGpp pools.

To test our *in vitro* model of (p)ppGpp–c-di-GMP cross-regulation, we turned to genetic approaches and *in vivo* analysis of c-di-GMP–dependent phenotypes. The degradation of CtrA, a master regulator of cell-cycle progression in *C. crescentus*, is tightly regulated by c-di-GMP levels. Briefly, c-di-GMP binding to the PopA adaptor drives CtrA degradation by the ClpXP protease at the swarmer-to-stalked transition (Duerig et al., 2009; Ozaki et al., 2014). Under favorable conditions, while PleD is not essential, its activation and the resulting c-di-GMP synthesis has been shown to promote both cell differentiation and CtrA degradation to drive cell-cycle progression (Fig. 5D) (Abel et al., 2013; Kaczmarczyk et al., 2020; Lori et al., 2015; Paul et al., 2007). Conversely, under glucose exhaustion conditions, wild-type cells maintain CtrA protein levels, which effectively blocks differentiation (Boutte et al., 2012)(Fig. 5D, Fig. S6A). This observation correlates with the hypothesis that (p)ppGpp accumulation in this condition inhibits PleD’s activity, preventing the rise in c-di-GMP levels required for CtrA degradation.

To test this model, we examined CtrA levels in a *rel* mutant, which does not produce (p)ppGpp. In a *rel* mutant, *i.e.* in the absence of (p)ppGpp, CtrA follows a typical cell-cycle dependent profile under favorable conditions, while under glucose exhaustion, CtrA is degraded (Fig. 5D). This suggests that, without (p)ppGpp, the burst in c-di-GMP synthesis occurs which triggers CtrA degradation. In support of that, inactivation of *pleD* in *rel* mutant cells suppressed the CtrA degradation under glucose exhaustion, with the *rel pleD* double mutant behaving like the wild-type and the *pleD* mutant (Fig. 5D, Fig. S6D). In contrast, CtrA oscillations were maintained under favorable conditions in both the *pleD* single mutant and the *rel pleD* double mutant (Fig. 5D; Fig. S6D). Notably, despite CtrA degradation under glucose exhaustion, *rel* mutant cells do not start DNA replication (Fig. S6A), likely due to repression of *dnaA* expression and rapid degradation of the protein under starvation conditions, as previously reported (Hallgren et al., 2023; Leslie et al., 2015).

Additionally, we compared the growth of wild-type, *rel* mutant, and *rel pleD* mutant strains under glucose exhaustion conditions. We found that in both asynchronous populations and synchronized cells, growth halted only in the wild-type and *rel pleD* mutant strains, while the *rel* mutant continued to grow slightly (Fig. S6EF). This observation aligns with our hypothesis that the accumulation of (p)ppGpp in wild-type cells under starvation conditions prevents cell-cycle transition to growth and differentiation by inhibiting PleD. Moreover, the fact that unsynchronized *rel* mutant cells grow more than one doubling time under glucose exhaustion conditions, whereas wild-type cells (and *rel pleD* mutant cells) stop after one doubling time (Fig. S6E), indicates that (p)ppGpp production is responsible for halting growth before glucose or other carbon sources are fully depleted, and suggests a PleD-dependent mechanism. Of note, this also explains our choice to resuspend synchronized cells in spent medium rather than M2 medium for experiments: spent medium retains residual glucose and carbon sources, allowing cells to continue progressing through the cell cycle without triggering abrupt starvation, as M2 medium would.

To further test that elevated c-di-GMP levels are responsible for the CtrA degradation observed in glucose exhausted *rel* mutant cells, we expressed the phosphodiesterase *proE* from *P. aeruginosa* in the *rel* mutant under glucose exhaustion conditions. Ectopic expression of *proE* has been shown to reduce c-di-GMP levels in C. crescentus (Duerig et al., 2009) and we observed that it restored CtrA stabilization under glucose exhaustion conditions whereas a catalytic mutant (*proE\**) could not (Fig. 5E; Fig. S6A), further supporting the idea that PleD-dependent c-di-GMP synthesis drives CtrA degradation in *rel* mutant cells in carbon limiting conditions.

## Discussion

Our results demonstrate that (p)ppGpp directly inhibits the diguanylate cyclase activity of PleD by competing with its GTP substrate at the enzyme’s active site. *In vitro*, both ppGpp and pppGpp bind PleD (Fig. 3A, Fig. S2) and significantly reduce c-di-GMP synthesis in a concentration-dependent manner (Fig. 3BC, Fig. S3). This inhibition occurs independently of PleD’s phosphorylation state (Fig. 3D, Fig. S3C), indicating that (p)ppGpp modulates PleD activity outside of its canonical activation pathway. Structural data further support a mechanism where (p)ppGpp binds within the GGDEF catalytic site and stabilizes an inactive dimeric conformation of PleD (Fig. 4, Fig. S4).

*In vivo*, pppGpp accumulates under carbon exhaustion while GTP levels decline (Fig. 5BC), consistent with a model in which pppGpp links nutrient limitation to cell-cycle control by suppressing PleD-dependent c-di-GMP synthesis. Under carbon exhaustion, pppGpp concentrations rise to ∼10 µM, which remains lower than ppGpp concentrations in the same conditions (Fig. 5AB) but remains in the same range as the K_D_ measured in the ITC assays we performed (Fig. 3A). We observed that (p)ppGpp concentrations in *C. crescentus* remain well below those typically reported in *E. coli* using quantitative LC–MS methods, where measured basal levels were ∼140 µM for ppGpp and ∼4 µM for pppGpp, and could rise to several hundred micromolar under stringent response (Patacq et al., 2018, 2020;Fig. 5B). This may reflect the oligotrophic lifestyle of *C. crescentus*, which may rely on a dampened alarmone response to cope with frequent nutrient variation. Supporting this model, and as previously published (Boutte et al., 2012; Hallgren et al., 2023), we observed a Rel-dependent stabilization of the master cell-cycle regulator CtrA under carbon exhaustion in synchronized cultures (Fig. 5D). Conversely, in *rel* mutant cells, which lack (p)ppGpp accumulation, CtrA was degraded (Fig. 5D). This phenotype was PleD- and c-di-GMP-dependent, as CtrA was stabilized in *rel pleD* double mutants and in *rel* mutant cells ectopically producing the c-di-GMP-degrading enzyme ProE (Fig. 5DE).

Our structural and biochemical analyses indicate that direct inhibition of DGCs by (p)ppGpp may be conserved across enzymes with shared architecture. In *C. crescentus*, (p)ppGpp binds within the active site of PleD, competing with GTP and stabilizing an inactive dimeric conformation (Fig. 4C). This mechanism involves a conformational shift where the α4 helices of the GGDEF domains are <5 Å apart in the (p)ppGpp-bound form, compared to ∼20 Å in the product-bound structure (Fig. 4C). This “closed” arrangement strongly supports an active and specific mechanism of inhibition, rather than simple competitive binding. Notably, residues mediating (p)ppGpp binding, including R300, are conserved across PleD-family DGCs (Fig. S5), suggesting that this regulatory interaction may be broadly conserved. In addition to R300, our structural analysis identifies Y439 and R446 as residues that specifically engage the γ-phosphate of pppGpp. All three residues are highly conserved features of GGDEF domains across DGCs (Fig. S5), and R300 and R446 have been shown to be important for catalytic function (Wassmann et al., 2007;Fig. S4D), contributing to both GTP binding and (p)ppGpp-mediated inhibition. This dual involvement suggests that mutations disrupting (p)ppGpp binding are likely to impair PleD activity itself, making it difficult to uncouple the inhibitory and catalytic functions.

Orthologs of PleD from *A. tumefaciens* and *S. meliloti* are similarly inhibited by (p)ppGpp *in vitro*, with pppGpp showing stronger effects, consistent with our findings in *C. crescentus* (Fig. 3BEF). These enzymes share a conserved domain organization, with two N-terminal receiver domains that promote dimerization upon phosphorylation and a C-terminal GGDEF domain responsible for catalysis. This conservation suggests that coupling DGC activity to metabolic stress via (p)ppGpp may be a widespread strategy among α-proteobacteria. WspR from *Pseudomonas aeruginosa* is a PleD ortholog that also adopts this two-domain architecture and can be activated by phosphorylation. However, unlike PleD, WspR can dimerize and become active independently of phosphorylation (De et al., 2008). We found that WspR is partially inhibited by pppGpp *in vitro*, suggesting that alarmone-mediated regulation may extend beyond α-proteobacteria, albeit with different sensitivities. By contrast, DgcZ from *E. coli*, which features a Zn-sensing regulatory domain and forms constitutive dimers (Zähringer et al., 2013), shows no significant activity change in the presence of (p)ppGpp *in vitro* (Fig. 3H). This highlights that GGDEF domain conservation alone is not sufficient to predict functional impact, which likely depends on additional factors such as enzymatic turnover rate, oligomerization dynamics or other mechanisms. These differences underscore the structural and kinetic complexity shaping how DGCs respond to intracellular alarmone signals. In the case of PleD, (p)ppGpp binding not only blocks substrate access but also locks the enzyme in an inactive conformation, providing a clear example of active inhibition. These findings argue against a purely competitive model and underscore the need to consider the structural and mechanistic diversity of (p)ppGpp regulation. Further studies will be needed to determine how widespread this mechanism is across bacterial DGC families and how it integrates with broader stress and signaling networks. Notably, when purifying PleD *in vitro*, we recovered a fraction of PleD with c-di-GMP bound, likely at its I-site, consistent with observations for other GGDEF enzymes (Chan et al., 2004; De et al., 2009; Wassmann et al., 2007). However, PleD_SM_ and DgcZ preparations contained little or no bound c-di-GMP (Fig. S2A). Of note, these DGCs show higher c-di-GMP synthesis activity, which is consistent with the inhibitory effect of c-di-GMP upon binding to the I-site (Fig. S3HI). However, the absence of pre-bound ligand in these enzymes did not increase their sensitivity to (p)ppGpp inhibition, indicating that the inhibitory effect is not simply due to displacement of I-site-bound c-di-GMP, but rather reflects a direct interference with catalytic activity.

Interestingly, despite the strong inhibitory effect of (p)ppGpp on PleD activity *in vitro*, we did not detect significant differences in c-di-GMP concentrations between nutrient-rich and starvation conditions at the population level (Fig. S6C). Similar measurements in synchronized populations have previously revealed clear cell-cycle oscillations of c-di-GMP, with levels rising during the swarmer-to-stalked transition (Abel et al., 2013). Thus, the absence of changes in our bulk assays may reflect averaging over heterogeneous subpopulations, masking temporal variations in c-di-GMP accumulation. The absence of major changes in c-di-GMP levels could also reflect compensatory activities or the activity of other diguanylate cyclases that remain functional under nutrient limitation, or alternatively, a spatial compartmentalization of c-di-GMP synthesis and degradation. Such subcellular organization of c-di-GMP pools has already been described in *C. crescentus* (Christen et al., 2010; Kaczmarczyk et al., 2020, 2024). In *C. crescentus*, (p)ppGpp levels may fluctuate across the cell cycle and may be higher in swarmer cells than in stalked cells (Ronneau et al., 2016; Shyp et al., 2020). At the same time, PleD is transiently recruited to the cell pole during the swarmer-to-stalked transition, where it triggers a burst of c-di-GMP synthesis (Abel et al., 2011; Christen et al., 2010; Paul et al., 2008). This spatiotemporal coordination raises the possibility that PleD activity is controlled by local or transient changes in the GTP:(p)ppGpp ratio, which may not be reflected in bulk measurements. While this hypothesis remains to be tested, it highlights the importance of considering spatial and temporal dynamics when interpreting second messenger signaling. Similar principles have been observed in other bacteria, where local control of nucleotide signaling enables precise regulation of specific outputs, independently of global nucleotide levels (Nicastro et al., 2020; Richter et al., 2020).

Beyond its direct inhibition of PleD, (p)ppGpp may also influence c-di-GMP signaling indirectly by reshaping nucleotide metabolism. As a central regulator of purine biosynthesis, (p)ppGpp lowers intracellular GTP levels during nutrient stress, reducing substrate availability for DGCs (Fig. 5C). Such interplay between purine metabolism and c-di-GMP signaling has been described in *P. aeruginosa* (Kennelly et al., 2024). Although we did not observe major changes in bulk c-di-GMP levels in *C. crescentus*, this dual-layer regulation, combining enzyme inhibition and precursor limitation, may contribute to the fine-tuning of local signaling.

While our proposed model is that (p)ppGpp directly inhibits PleD to modulate cell differentiation, this is likely only one facet of its broader regulatory role in *C. crescentus*. As in other bacteria, (p)ppGpp likely influences multiple core processes, including transcription, translation, and metabolism in *C. crescentus*. Thus, while the PleD pathway provides a clear example of how metabolic signals are coupled to developmental outcomes, it likely operates in parallel with additional layers of (p)ppGpp-dependent regulation that collectively shape the growth and cell cycle dynamics of *C. crescentus*.

The interaction between (p)ppGpp and DGCs such as PleD adds to a growing body of evidence that alarmone signaling intersects with c-di-GMP pathways at multiple levels. In addition to modulating DGC activity, (p)ppGpp also regulates effector proteins like SmbA, which has been shown to bind both (p)ppGpp and c-di-GMP at the same site to control growth in *C. crescentus* through a metabolic switch(Shyp et al., 2020). This work underscores the potency of second messengers to function as molecular switches at the interface between metabolism and development, and highlights how direct cross-talk between signaling pathways can coordinate complex cellular decisions in bacteria.

## Acknowledgments

We acknowledge the contribution of SFR Biosciences (Université Claude Bernard Lyon 1, CNRS UAR3444, Inserm US8, ENS de Lyon) at the Protein Science Facility, especially Eric Diesis and Virginie Gueguen-Chaignon. We acknowledge the contributions of the CELPHEDIA Infrastructure (http://www.celphedia.eu/), especially Sébastien Dussurgey at the center AniRA in Lyon. We thank Proxima 2 beamline staff scientists from SOLEIL and Dr. Max Nanao from beamline ID23EH2 of the ESRF for assistance during data collection. We are grateful to Michael Laub’s lab and the Koch Institute Biopolymer & Proteomics Core at MIT for helping us with the chemical proteomics work. We thank the Jault and Lesterlin labs at MMSB for technical support and access to equipment, especially Kevin Lang, Elise Kaplan, Cécile Gonzalez and Annick Dedieu, as well as Marie-Laure Fogeron for assistance with ITC. This research was supported by an ATIP Avenir grant funded by the Centre National de la Recherche Scientifique (CNRS) and the Fondation Bettencourt-Schueller awarded to MG, by an Amorçage Europe grant of the Région Auvergne-Rhône-Alpes awarded to MG and by a FEBS Excellence Award from the Federation of European Biochemical Societies awarded to MG. We thank Michael Laub and Justine Collier for critical reading of the manuscript.

## Contributions

Conceptualization: C.J and M.G - Methodology: C.J, C.D, J.P-S.M, P.V, F.L, R.H, B.W, L.T, M.G - Formal analysis: C.J, C.D, J.P-S.M, P.V, F.L, R.H, B.W, L.T, M.G - Investigation: C.J, C.D, J.P-S.M, P.V, B.W, L.T, M.G - Writing: original draft from C.J and M.G with contributions of all authors-Resources: L.T., R.H., F. L. and M.G. - Funding acquisition: M.G

## Declaration of interests

The authors declare no competing interests.

## Methods

### Growth conditions

*Caulobacter crescentus* strains were grown in PYE (2 g/L bactopeptone, 1g/L yeast extract, 0.3 g/L MgSO_4_, 0.5 mM CaCl_2_) or in fresh M2 medium (0.87 g/L Na_2_HPO_4_, 0.53 g/L KH2PO_4_, 0.5 g/L NH_4_Cl, 0.5 mM MgSO_4_, 10 mM FeSO_4_, 0.1 mM CaCl_2_) supplemented with 0.2% or 0.02% glucose at 30°C with constant shaking (210 rpm). Expression from the P_van_ promoter was induced with vanillate (100 µM). To maintain plasmids, antibiotics were added at the following concentrations (liquid/plate): kanamycin (5 μg mL^−1^ / 25 μg mL^−1^), oxytetracycline (1 μg mL^−1^ / 2 μg mL^−1^), gentamycin (2.5 μg mL^−1^ / 5 μg mL^−1^), chloramphenicol (2 μg mL^−1^ / 1 μg mL^−1^).

*Escherichia coli* strains were grown in LB (10 g/L NaCl, 10 g/L tryptone, 5 g/L yeast extract) and supplemented with antibiotics at the following concentrations unless noted (liquid/plate): kanamycin (30 μg mL^−1^ / 50 μg mL^−1^), oxytetracycline (12 μg mL^−1^ /12 μg mL^−1^), gentamycin (15 μg mL^−1^ / 20 μg mL^−1^), chloramphenicol (20 μg mL^−1^ / 30 μg mL^−1^), ampicillin (100 µg mL^−1^ /100 µg mL^−1^).

### Strain construction

*Escherichia coli* strains were derivative of the BL21(DE3) strain for the production of recombinant proteins. All BL21 derivative strains were constructed by transforming the replicative plasmid by heat-shock. All *C. crescentus* strains were derivative of the wild-type isolate MGC001 from the CB15N/NA1000 strain. All strains, plasmids and oligonucleotides used in this study are listed in Tables S5-S7.

To construct strains MGC015 and MGC016 and strains MGC564 and MGC565, MGC001 and MGC533 were electroporated with pBVMCS-4-*proE* or pBVMCS-4-*proE\** respectively, selecting on gentamycin plates.

Strain MGC636 (NA1000 *Δrel ΔpleD*) was constructed using two-step recombination [Skerker JM, Prasol MS, Perchuk BS, Biondi EG, Laub MT (2005)]. First, pNPTS138-*pleD::tet* was electroporated into MGC533 (NA1000 *Δrel*) followed by selection on kanamycin. Single colonies obtained were grown overnight in liquid PYE + tetracycline. After 10-15 hour-growth, 5 μL in 45 μL PYE was plated for counter-selection on PYE containing tetracycline and 0.3% sucrose. Sucrose resistant clones were isolated on PYE + tetracycline and PYE + tetracycline + kanamycin plates and isolated clones that lost kanamycin resistance were selected. *pleD* deletion was confirmed by PCR.

### Plasmid construction

pNTPS138-*pleD*: The *pleD* deletion cassette was amplified from MGE477 chromosomal DNA using primers MGP841 and MGP842. Fragment was cloned into EcoRV site of the pNTS138 plasmid using DNA ligation. The construction was verified by sequencing.

pET28a-*divJ_1-246_:* The *divJ_1-246_* fragment encoding DivJ deleted of the first 246 amino acids was amplified from *Caulobacter crescentus* NA1000 chromosomal DNA using primers MGP817 and MGP821. Fragment was cloned into BamHI restriction site of the pET28a plasmid using <u>S</u>equence and <u>L</u>igation Independent <u>C</u>loning (SLIC). The construction was verified by sequencing.

pET28a-*pleD_AT_*: The *pleD_AT_* fragment was amplified from *Agrobacterium tumefaciens* C58 chromosomal DNA using MGP856 and MGP857. The pET28a plasmid was amplified using MGP730 and MGP855. Fragment was cloned into amplified pET28a using SLIC. The construction was verified by sequencing.

pET28a-*pleD_SM_*: The *pleD_SM_* fragment was amplified from *Sinorhizobium meliloti* 1021 chromosomal DNA using MGP899 and MGP900. The pET28a plasmid was amplified using MGP730 and MGP855. Fragment was cloned into amplified pET28a using SLIC. The construction was verified by sequencing.

pET28a-*dgcZ*: The *dgcZ* fragment was amplified from *Escherichia coli* MG1655 chromosomal DNA using MGP 858 and MGP 859. The pET28a plasmid was amplified using MGP730 and MGP855. Fragment was cloned into amplified pET28a using SLIC. The construction was verified by sequencing.

pET28a-*wspR:* The *wspR* fragment was amplified from *Pseudomonas aeruginosa* strain PA01 chromosomal DNA using MGP 860 and MGP 861. The pET28a plasmid was amplified using MGP730 and MGP855. Fragment was cloned into amplified pET28a using SLIC. The construction was verified by sequencing.

pET11b-*pleD_R300A_:* The pET11b-*pleD* plasmid was amplified using MGP1013 and MGP1014. The synthetic gBlock *pleD_R300A_* (Table S7) was cloned into amplified pET11b-*pleD* using SLIC. The construction was verified by sequencing.

pET11b-*pleD_R300K_*: The pET11b-*pleD* plasmid was amplified using MGP1013 and MGP1014. The synthetic gBlock *pleD_R300K_* (Table S7) was cloned into amplified pET11b-*pleD* using SLIC. The construction was verified by sequencing.

### Solid-phase peptide synthesis

We synthesized four peptides with the following sequence: biotinyl-Gly-Lys(bromoacetyl-X-pMet-)-Gly_-_NH_2_(Wang et al., 2019). X is a linker amino acid varying in each peptide.

| Peptide # | X | Free amino-acid structure for X |
| --- | --- | --- |
| 1 | Gly | H<sub>2</sub>N-CH<sub>2</sub>COOH |
| 2 | PEG2 | H<sub>2</sub>N-(CH<sub>2</sub>CH<sub>2</sub>O)<sub>2</sub>-CH<sub>2</sub>COOH |
| 3 | PEG4 | H<sub>2</sub>N-(CH<sub>2</sub>CH<sub>2</sub>O)<sub>4</sub>-CH<sub>2</sub>COOH |
| 4 | PEG6 | H<sub>2</sub>N-(CH<sub>2</sub>CH<sub>2</sub>O)<sub>6</sub>-CH<sub>2</sub>CH<sub>2</sub>COOH |

The general procedure for a coupling-deprotection cycle: 5 eq. Fmoc protected amino acid and 4.8 eq. of 1-[Bis(dimethylamino)methylene]-1H-1,2,3-triazolo[4,5-b] pyridinium 3-oxide hexafluorophosphate (HATU) dissolved in N,N-dimethylformamide (DMF) to achieve a final concentration of 0.4 M amino acid and 0.38M HATU. 5 eq. of diisopropylethylamine (DIEA) was then added and the mixture was applied to the solid support and left to react at room temperature (RT) for 20 minutes, followed by 3 washes with DMF, 2 x 8 min deprotection with 20% (v/v) piperidine in DMF and 5 washes with DMF. Biotin was coupled following the same procedure but without deprotection. For coupling of Fmoc-pMet-OH, usage of the amino acid, HATU and DIEA were reduced to 2.0, 1.9, and 2 eq., respectively, and coupling time extended to 1 hour at RT.

The Biotinyl-GKG structure first synthesized at 0.1-mmol scale on Rink-amide resin (ChemMatrix). The resin was swelled in DMF prior to the first coupling. Coupling-deprotection cycles were performed with Fmoc-Gly-OH, Fmoc-Lys(Alloc)-OH, and Fmoc-Gly-OH (in this order), and the N-terminal amine was biotinylated. Thereafter, The Alloc protecting group was quantitatively removed by a 20-minute, RT treatment with 598 µL of phenylsilane (4.85 mmol) and 45.63 mg (39.5 µmol) of tetrakis(triphenylphosphine)palladium(0) in 4 mL of dichloromethane (DCM). The resin was then thoroughly washed with dichloromethane (DCM), air-dried, and stored in a desiccator for further use.

Syntheses was continued on a 0.01-mmol scale. Resin bearing the Biotinyl-GKG fragment was weighed into individual vessels and re-swelled with DMF. Fmoc-pMet-OH (Prepared in-house from L-Photo-Methionine, Thermo Fisher) was coupled to the lysine side chain followed by the linker amino acid X: Fmoc-Gly-OH (Sigma-Aldrich), Fmoc-NH-PEG2-CH_2_COOH, Fmoc-NH-PEG4-CH_2_COOH or Fmoc-NH-PEG6-CH_2_CH_2_COOH (ChemPep). The deprotected N-terminus of the linker residue was then bromoacetylated by 500 µL DMF solution containing 25 mg bromoacetic anhydride (10 eq) and 3.5 µL DIEA (2 eq) for10-minute incubation at RT. Finally, the resin bearing each peptide was washed thoroughly with DCM, air dried, and the product peptide was cleaved from the resin with 1 mL neat trifluoroacetic acid (TFA) containing 2.5% (v/v) water and 2.5% (v/v) triisoproprylsilane at 60 °C for 8 min. TFA in the resulting solution was evaporated and the residual was triturated with 1+1 (v/v) mixture of diethyl ether and hexane to precipitate the peptide. The crude peptide was washed once with the ether-hexane mixture, air-dried and lyophilized. The crude peptide was refined over a semi-preparative reverse phase (RP)-HPLC (Discovery BIO wide pore C18 column: 10 x 250 mm, 5 µm) using a linear gradient of solvent A (0.1% TFA in 90% acetonitrile) and B (0.1% TFA) with the percentage of B increasing from 5% to 25% over 10 minutes. Flow rate was 6 mL/min. Pure fractions of the product peptide were combined and lyophilized.

### Synthesis of ppGpp photo-affinity probes

Purified *B. subtilis* SasB was used to synthesize [5’-β-thio]-ppGpp (ppGpp-5βS) and [3’-β-thio]-ppGpp (ppGpp-3βS). The reactions were conducted in 2mL buffer containing 20mM Tris-HCl 8.0, 150 mM NaCl, 25mM MgCl2 and 1mM TCEP. Final concentrations were 12 mM ATP and 10 mM [5’-β-thio]-GDP (Sigma-Aldrich) for ppGpp-5βS synthesis and 10 mM [5’-β-thio]-ATP (Sigma-Aldrich) and 10 mM GDP for ppGpp-3βS synthesis. 250 µM pppGpp was included to activate SasB(Steinchen et al., 2015). This mixture was incubated with 5 µM SasB at 37°C for 3 hours and completion of reaction confirmed by anion-exchange (ANIEX) chromatography on a MonoQ 5/50 column using buffers A: 5 mM Tris-HCl 8.0 and B: 5mM Tris-HCl 8.0 and 1mM NaCl. A linear gradient of 10-40%B in 15 mL was used. All ANIEX runs were completed with Upon completion, the reaction was vortexed with 300uL chloroform for 1 minute, and spun at 21,000 *g* for 5 min. The aqueous layer was diluted 20-fold in water and resolved on MonoQ 10/100 with 10-40%B in 80mL. Fractions containing the product were combined, lyophilized and re-dissolved in 5 mL water. LiCl powder was then added to 1 M concentration, and the products were precipitated by addition of 20 mL ice-cold 100% ethanol. After staying in ice-water bath for 30 min, precipitate was spun down, mother liquor aspirated, and pellet was washed with 5 mL 95% EtOH. The final product was collected through centrifugation, redissolved in water, quantified based on absorption at 252 nm (ε = 13,600 M^−1^ cm^−1^) and then lyophilized. Dried, colorless powder was dissolved with Milli-Q water for 100mM stocks and stored frozen −80°C. ppGpp photo-affinity probes were synthesized by conjugating 2.2 µmol ppGpp-5βS or ppGpp-3βS to 2 µmol peptide at pH = 6.5 at 42 °C for 1 hr. Conjugation to peptide **1**-**4** were carried out in separate reactions. Products were purified on a MonoQ (5/50) column at 4 °C using buffer A (5 mM HEPES-Na 7.0, 10% acetonitrile) and buffer B (5 mM HEPES-Na pH 7.0, 1M NaCl, 10% acetonitrile). All Four ppGpp-5βS conjugates were then combined into an equimolar mixture and diluted to 0.5 mM total concentration. A similar stock was prepared for all four ppGpp-3βS conjugates.

Effector capture and enrichment from *C. crescentus* lysate

M2G medium containing 0.4% glucose and 0.50 g/L regular NH_4_Cl (light medium) or 0.51 g/L NH_4_Cl (^15^N, 99%, Cambridge Isotope Laboratories, heavy medium) were used. *C. crescentus NA1000* was cultured at 30°C in 60-mL batches in 500-mL baffled flask with vigorous shaking to OD_600_ = 0.38 and then chilled on ice. Cells were collected, washed with 1 mL lysis buffer (20 mM HEPES-Na 7.0, 300 mM NaCl), and resuspended in lysis buffer to 600 µL total volume. This sample was sonicated and cell debris removed through centrifugation at 20,000G for 10 minutes to yield a cleared lysate containing approximately 6 mg/mL total protein. Two reactions were assembled in adjacent wells on a 96-well plate chilled on ice for each photo-affinity probe mixture. The effector-capture reaction consisted of 400 µg protein from the light lysate, 100 µM photo-affinity probe, 1.8 mM MgCl_2_, and 0.2 mM MnCl_2_. The control reaction consisted of 800 µg protein from the heavy lysate, 100 µM capture compound, 5 mM ppGpp, 6.3 mM MgCl_2_ and 0.2 mM MnCl_2_. After exposure to 365-nm UV lamp for 2 minutes, reactions were sampled for western blotting and all four reactions (one pair each for ppGpp-5βS and ppGpp-3βS probes) were combined and extensively exchanged in a 10kDa-MWCO concentrator into a buffer containing 20 mM HEPES-Na pH 7.5 and 200 mM NaCl. The final concentrate (approximately 100 µL) was diluted with 400 µL RIPA buffer (50 mM Tris 7.5, 150 mM NaCl, 0.1% SDS, 0.5% sodium deoxycholate, 1% Triton X-100) and incubated overnight with 200 µL MyOne Streptavidin C1 dynabeads (Thermo Fisher Scientific) at 4 °C. On the second day, the protein in supernatant (flowthrough) was sampled for Western blotting and then removed, and the dynabeads were washed at 4 °C twice with RIPA buffer, once with 0.1 M Na_2_CO_3_, once with 100 mM Tris 8.0, 4 M guanidinium chloride (GuHCl) and twice more with RIPA buffer. Bound protein was eluted through boiling the dynabeads in 40 µL SDS-PAGE loading dye containing 2 mM biotin. The eluate was lyophilized and re-diluted using water to 15 µL, resolved on 12% TGX-PAGE.

### LC-MS2 based proteomics

SDS-PAGE of eluates from streptavidin dynabeads was stained with Coomassie Brilliant Blue (CBB), revealing three major protein bands, the streptavidin at 13 kDa, EF-Tu at 40 kDa and EF-G at 80 kDa. The gel lane for each eluate was excised and regions below the 15-kDa or above the 200-kDa marker were trimmed off. The lane was further divided into three samples and processed separately: the EF-Tu and EF-G bands (sample A), the fragment below EF-Tu (sample B), and the rest (sample C). The gel fragments were diced into 1-mm pieces and then treated with 10 mM DTT at 56 °C for 1 hr followed by 55 mM iodoacetamide at RT for 1 hr in the dark. After a brief wash with 0.1 M NH_4_HCO_3_, gel pieces were dehydrated with acetonitrile, dried in a speedvac, and re-hydrated with a minimal volume of 6 ng/µL trypsin in 0.1 M NH_4_HCO_3_ for digestion at RT overnight. Thereafter, tryptic fragments were extracted by four shrinking-swelling iterations. In each iteration, gel pieces were first dehydrated with 5% formic acid in 50% (v/v) acetonitrile (first two iterations) or neat acetonitrile (second two). Supernatant containing tryptic fragments was then removed for collection and gel pieces were re-swelled with 0.1 M NH_4_HCO_3_ in water. The combined supernatant was dried in a speedvac and reconstituted in 50 µL 0.1% formic acid. 5 µL was used for LC-MS2 analysis.

Tryptic fragments were separated by RP-HPLC (Thermo Easy nLC1000) using a precolumn (made in house, 6 cm of 10 µm C18) and a self-pack 5 µm tip analytical column (12 cm of 5 µm C18, New Objective) over a 140-minute gradient before nanoelectrospray using a QExactive mass spectrometer (Thermo). Raw mass spectral data files (.raw) were searched using Proteome Discoverer 2.1 (Thermo) and Mascot version 2.4.1 (Matrix Science). Mascot search parameters were: 10 ppm mass tolerance for precursor ions; 15 mmu for fragment ion mass tolerance; 2 missed cleavages of trypsin; fixed modification was carbamidomethylation of cysteine. A separate search was carried out for heavy peptides whereby replacement of all ^14^N by ^15^N in all amino acids were included as fixed modification. Signals of heavy and light fragments of the same sequence are related by expected mass difference and retention time. Quantification was then obtained using the area of the precursor-ion peaks. SILAC ratio of a unique peptide sequence was calculated as the mean of all precursor ions that match the sequence regardless of modifications. SILAC ratio of a protein was calculated as the mean of all unique peptides mapped to the protein. SILAC ratios were converted to log_2_ units. Only proteins detected in both replicates each by at least two unique tryptic fragments were reported.

### Purification of His-tagged proteins

All Histidine-tagged proteins were purified with the standard protocol described below. *E. coli* BL21(DE3) were used to express proteins from pET11b and pET28a expression plasmids (Table S5, S6). Cells were grown in 1-4 L of LB medium supplemented with ampicillin for cells carrying pET11b-derived plasmid or kanamycin (25 µg/mL) for cells carrying pET28a-derived plasmid. Expression was induced by adding isopropyl 1-thio-D-galactopyranoside (IPTG) to 0.5 mM final concentration and the cultures were incubated overnight at 20°C. After harvesting by centrifugation, the cell pellets were resuspended in 40 mL lysis buffer containing 50 mM Tris-HCl, pH 8.0, 500 mM NaCl, 10 mM imidazole, 5 mM β-mercaptoethanol, 50 µg mL^−1^ lysozyme (Sigma), 5 µg/mL DNase I (Roche) and Complete protease inhibitor (Roche). Cells were disrupted through sonication, and the lysates were cleared at 100,000 x *g* for 1 hr. at 4°C. Cleared lysate was applied to 5 mL Ni–NTA resin equilibrated with wash buffer 1 (50 mM Tris-HCl, pH 8.0, 500 mM NaCl, 10 mM imidazole, 5 mM β-mercaptoethanol) on an ÄKTA pure™ system (GE Healthcare). The Ni– NTA resin was washed with ten column volumes of wash buffer 1. The bound protein was eluted with a linear gradient of elution buffer (50 mM Tris–HCl pH 8, 500 mM NaCl, 5 mM β-mercaptoethanol and 400 mM imidazole). Elution fractions were pooled and dialyzed overnight in SEC buffer (25 mM Tris–HCl pH 7.8, 100 mM NaCl, 5 mM MgCl_2_, glycerol 5% and 1 mM β-mercaptoethanol). Dialyzed proteins were concentrated using a 30kDa Amicon® Ultra Centrifugal concentrator (Merck Millipore). Then, the concentrated proteins were centrifuged at 16,000 x *g* for 15 min. at 4 °C and loaded onto a Superdex™ Increase 200 (10/300) gel filtration column (GE Healthcare) equilibrated in SEC buffer. The fractions were pooled and concentrated to the desired concentration (around 10 mg ml^−1^) for further experiments and frozen at −80°C.

### Isothermal Titration Calorimetry (ITC)

Experiments were carried out at 25 °C, a syringe stirring speed of 700 r.p.m., a pre-injection delay of 60 sec. and a recording interval of 150 sec. in an iTC200 MicroCalorimeter (Malvern Panalytical). SEC purified PleD at 100 µM was loaded in the sample and the titration of the protein performed in SEC buffer (25 mM Tris–HCl pH 7.8, 100 mM NaCl, 5 mM MgCl_2_, glycerol 5% and 1 mM β-mercaptoethanol). ppGpp, pppGpp or GpCpp (Jena Biosciences) in SEC buffer were injected in the sample cell. Blank titrations were performed with protein-free SEC buffer in the sample cell. Baseline correction and integration of the raw differential power data and fitting of the resulting binding isotherms to obtain the K_D_ values were performed using the Microcal ORIGIN software. The parameters and the concentration of protein and ligands used are described in Table S1.

### nano Differential Scanning Fluorimetry (nanoDSF)

For nanoDSF experiments, 100 μM purified protein in 25mM Tris pH 7.8, 100mM NaCl, 5% (v/v) glycerol, 5mM MgCl_2_ and 1 mM β-mercaptoethanol was incubated without or with 1 mM ATP, GDP (Sigma), ppGpp, pppGpp, or GTP (Jena Biosciences), respectively, at room temperature for 5 min. Reactions were subsequently analyzed in a Prometheus NT.48 (NanoTemper Technologies) device with a 1°C/min. thermal ramp (from 20°C to 95°C) and the internal fluorescences at 330 nm and 350 nm were recorded. Data analysis and calculation of derivatives were done using the PR.ThermControl software (NanoTemper Technologies).

### Enzymatic assays

Diguanylate cyclase assays were adapted from procedures described previously (Paul et al, 2004). The reaction mixtures contained purified PleD_CC_, PleD_AT_, PleD_SM_, WpsR or DgcZ (40 µM) in 25 mM Tris-HCl pH 7.8, 100 mM NaCl, 10 mM MgCl_2_ in a 50 μL final volume. The reactions were started by the addition of 100 µM GTP to the mixture. For enzymatic assays of phosphorylated PleD, the protein was preincubated for 25 min. with purified DivJ_1-246_ (10 µM) and 200 µM ATP in the same buffer previously described. When added, the (p)ppGpp, ATP, GDP or GpCpp molecules (Jena Biosciences) were supplemented at the indicated concentration at the same time as the GTP substrate and incubated at 25°C for 2 hours. The reactions were stopped by addition of EDTA at a final concentration of 10 mM and transferred into the HPLC system (UltiMate3000, Thermo Fischer) for separation on a Nucleosil C18 column (Macherey-Nagel) with a 10 min. gradient of 70% acetonitrile – 0.09% trifluoroacetic acid. The c-di-GMP peak was detected using the 253 nm UV absorption.

### Crystallization, data collection, and structure determination

The screening was conducted using a Mosquito workstation (TTP Labtech) on commercial crystallization solutions with the sitting-drop vapor diffusion method. Drop were set up at 19°C and consisted of 150 nL of reservoir solution and 150 nL of protein solution (5 mg/mL) supplemented with 0.5 µM c-di-GMP and 5 mM ppGpp or pppGpp. All crystallization trials were visualized on RockImager 182 (Formulatrix) and crystals appeared after one month. Crystal of PleD in complex with ppGpp and c-di-GMP grew in 0.1 M NaCl, 50 mM sodium cacodylate pH 6.5, 10% w/v PEG 4000 and 500 µM spermine (Natrix, Hampton research, E7 condition). Crystal of PleD in complex with pppGpp and c-di-GMP were obtained in 0.2 M Lithium citrate tribasic tetrahydrate, 20% w/v Polyethylene glycol 3.350 (PEG ION HT (Hampton research), D9 condition). PleD-ppGpp crystals were frozen in a reservoir solution supplemented by 15% Glycerol, and PleD-pppGpp crystals were frozen in a reservoir solution supplemented by 15% Glycerol and 5mM pppGpp. X-ray Data for PleD-(p)ppGpp crystals were collected at the ID23EH2 beamline of the European synchrotron Radiation Facility and at Proxima 2 beamline of synchrotron SOLEIL. PleD-ppGpp crystals diffracted at a resolution 2.9 Å and PleD-pppGpp diffracted at 3 Å and both belonged to the space group P 6_1_22 with 4 molecules per asymmetric unit. Diffraction data were processed using XDS ((Kabsch, 2010) and with AIMLESS (Evans & Murshudov, 2013) from the CCP4 program suite (Hough & Wilson, 2018; “The CCP4 Suite: Programs for Protein Crystallography,” 1994). Data collection statistics are indicated in supplementary data Table S3.

Both the PleD-ppGpp and pppGpp structures were solved by molecular replacement using the coordinates of PleD monomer in complex with c-di-GMP (PDB ID: 1W25, (Chan et al., 2004)) as a probe in PHASER (Hough & Wilson, 2018). Models were re-built manually in COOT (Casañal et al., 2020) and refined using REFMAC (Murshudov et al., 2011) and PHENIX (van Zundert et al., 2021). The models were refined with final R_work_/R_free_ of 0.20/0.24 and 0.19/0.23 for PleD-ppGpp and PleD-pppGpp, respectively. The coordinates and structure factors were deposited in the Protein Data Bank with accession code 9G85 (PleD-ppGpp) and 9G86 (PleD-pppGpp). Figures were generated with Pymol, molecular Graphic System, Version 3 Schrödinger, LLC.

### Time courses and synchronization

Cells were grown overnight in PYE with appropriate antibiotics. Cells were diluted in M2 supplemented with 0.2% glucose and appropriate antibiotics when necessary and grown to OD_600_ 0.1-0.4. Cells were either diluted in M2 supplemented with 0.2% glucose to an OD_600_ of 0.06 or switched to M2 supplemented with 0.02% glucose (M2G_1/10_) (for glucose exhaustion) to an OD_600_ of 0.13 and grown for 5 hours to both reach an OD_600_ of ∼0,35.

For synchronization, cells were first centrifuged at 10,000 × *g* for 10 minutes at 4°C. Swarmer cells were then isolated using a Percoll density gradient centrifugation. The cell pellets were resuspended in equal volumes of cold M2 buffer (0.87 g/L Na₂HPO₄, 0.53 g/L KH₂PO₄, 0.5 g/L NH₄Cl) and Percoll (GE Healthcare), followed by centrifugation at 10,000 × *g* for 20 minutes. The upper ring was removed, while the lower ring, containing swarmer cells, was transferred to a new 15-mL Falcon tube. The swarmer cells were washed with 13 mL of cold M2 buffer and centrifuged at 10,000 × *g* for 5 minutes at 4°C. Finally, the pellet was resuspended in 2 mL of cold M2 buffer and subjected to a final centrifugation at 21,000 × *g* for 1 minute.

Synchronized swarmer cells were released into either M2 medium supplemented with appropriate antibiotics and 0.2% glucose (M2G condition) or into filtered M2G 0.02% glucose spent medium used during the 5-hour exhaustion step before synchronization (M2G_1/10_*). OD_600_ was adjusted to 0.1-0.2 for each condition. When appropriate, cultures were supplemented with inducers. At the indicated time points, samples were harvested from each flask for flow cytometry (0.15 ml) or immunoblotting (2 ml). For flow cytometry, samples were stored in 30% ethanol at 4°C. For immunoblotting, cells were centrifuged for 1 min. at 14,000g, supernatant was aspirated, and frozen in liquid nitrogen.

### Immunoblotting

Frozen pellets were resuspended on ice in 40 μl Laemmli sample buffer (2% [w/v] SDS, 10% glycerol, 60 mM Tris-Cl [pH 6.8], 0.01% [w/v] bromophenol blue, 1% β-mercaptoethanol) per OD_600_ unit and denatured at 95°C for 10 min. Samples were separated by 12% SDS-PAGE for 60 min. at 130 V at room temperature. Blocking was done in TBST buffer (TBS containing 0.1% Tween-20) with 5% (w/v) non-fat milk powder and antibodies were used at the following concentrations: anti-RpoA (1:7,500) (Bio-Legend), anti-CtrA (1:5,000), anti-FLAG (1:7500) (Merck) diluted in TBST 5% non-fat milk. Horseradish peroxidase (HRP)-conjugated secondary antibodies (ThermoFisher) were used at the following concentrations: anti-mouse (1:10,000) for anti-RpoA and anti-Flag immunoblotting and anti-rabbit (1:10,000) for anti-CtrA immunoblotting. The membranes were developed with SuperSignal™ West Pico PLUS Chemiluminescent Substrate (ThermoFisher) and visualized with a Fusion FX (Vilber). Band intensities were quantified using ImageJ software.

### Flow cytometry

A fraction of fixed cells from the time course sampling (corresponding to an OD_600_ of ∼0.005) were centrifuged at 6,000 rpm for 4 min. Pelleted cells were resuspended in 1 mL of Na_2_CO_3_ buffer containing 3 mg ml^−1^ RNase A (Qiagen) and incubated at 50°C for at least 4 hours or overnight to digest RNA.

Diluted samples were stained with 0.25 μM SYTOX Green (Invitrogen) for flow cytometric quantification of cellular DNA content. Cells were run on a Attune NxT (Invitrogen) (FSC = 360, SSC = 460, FITC = 320; Threshold: FITC > 1000) / MASCQuant (Miltenyi) (param: FSC = 290, SSC = 574, FITC = 458; Threshold: FITC > 1000) and 50,000 total SYTOX positive events were collected for each experiment.

### Sampling and Intracellular Metabolite Extraction

Cells were cultured until the desired growth phase (OD_660_ = 0.4), then harvested by withdrawing an appropriate volume corresponding to 8 x 10^7^ cells. The collected samples were immediately quenched by mixing with 5 mL of a precooled methanol:acetonitrile:water solution (4:4:2, v/v/v) at −20°C to rapidly stop metabolic activity (Millard et al., 2014). An internal standard consisting of 50 µL of a uniformly ^13^C-labeled *E. coli* metabolite extract (prepared as described in Patacq et al., 2018) was added to each sample (Millard et al., 2014). Samples were vortexed vigorously and incubated at −20°C for 2 hours to ensure metabolite extraction. Cell debris was removed by centrifugation at 16,000 × g for 5 minutes at 4°C, and the supernatants were dried overnight using a SpeedVac concentrator (Eppendorf Concentrator plus, Eppendorf AG, Hamburg, Germany). Dried extracts were stored at −80°C until analysis. Prior to analysis, samples were reconstituted in 250 µL of ultrapure water and analyzed using ion chromatography (IC; Thermo Scientific Dionex ICS-5000+, Dionex, Sunnyvale, CA, USA) coupled to a Q Exactive Orbitrap mass spectrometer (Thermo Fisher Scientific, Waltham, MA, USA) equipped with a heated electrospray ionization (HESI) source. Separation was achieved with a KOH gradient as follows: 7 mM (0–1 min), linear increase to 15 mM (1–9.5 min), held at 15 mM (9.5–20 min), ramped to 45 mM (20–30 min), then to 70 mM (30–33 min), followed by 100 mM (33.1–42 min), and re-equilibrated at 7 mM (42.5–50 min). The total run time was 50 minutes, and 10 µL were injected. Mass spectra were acquired in negative ion mode using full-scan acquisition at a resolution of 70,000 (m/z 200), with the following source parameters: capillary temperature 300°C, heater temperature 450°C, sheath gas flow 40 AU, auxiliary gas flow 20 AU, S-lens RF level 60%, and spray voltage −3.5 kV. Metabolites were identified based on exact m/z values with a 5 ppm mass tolerance, using Xcalibur software (Thermo Fisher Scientific) and processed with El-Maven (version 0.12.0, (Agrawal et al., 2019)). Four independent biological replicates were analyzed.

### Intracellular Metabolite Concentrations

First, metabolite concentrations in extracts were determined using calibration curves and the ratio of ¹²C- to ¹³C-labeled peak areas for GDP, GTP and (p)ppGpp; c-di-GMP was quantified using the ^12^C-labeled form only due to absence of ^13^C-labeled isotopologue in the ^13^C-labeled *E. coli* extract. Then, total cell volume was estimated from the cell counts on OD_660_ (1 OD = 2 × 10^9^ CFU/mL) and an average *C. crescentus* cell volume of 6.46 × 10⁻¹⁶ L (Snyder et al., 2020). Finally, intracellular concentrations were calculated by dividing metabolite concentration in extracts by the total cell volume.

---

**Figure S1 related to Figure 2:**
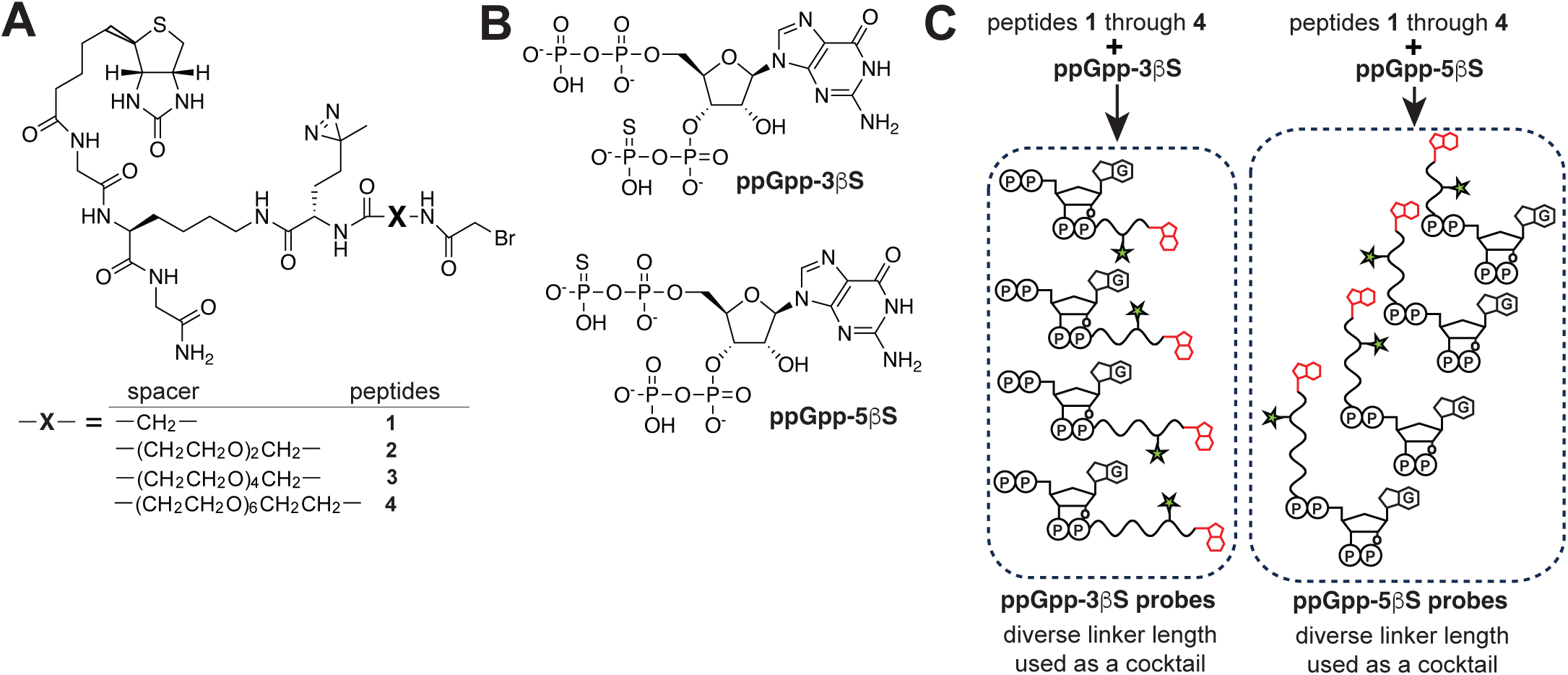
Synthesis of photo-affinity ppGpp probes^1^. Chemical structures of (A) bromoacetylated peptides containing a diazirine photocrosslinker and a biotin handle and (B) thiophosphate analogs of ppGpp are shown. (C) Conjugation of building blocks in (A) and (B) yields the tripartite photoaffinity probes used in Figure 2A-B.

**Figure S2 related to Figure 3:**
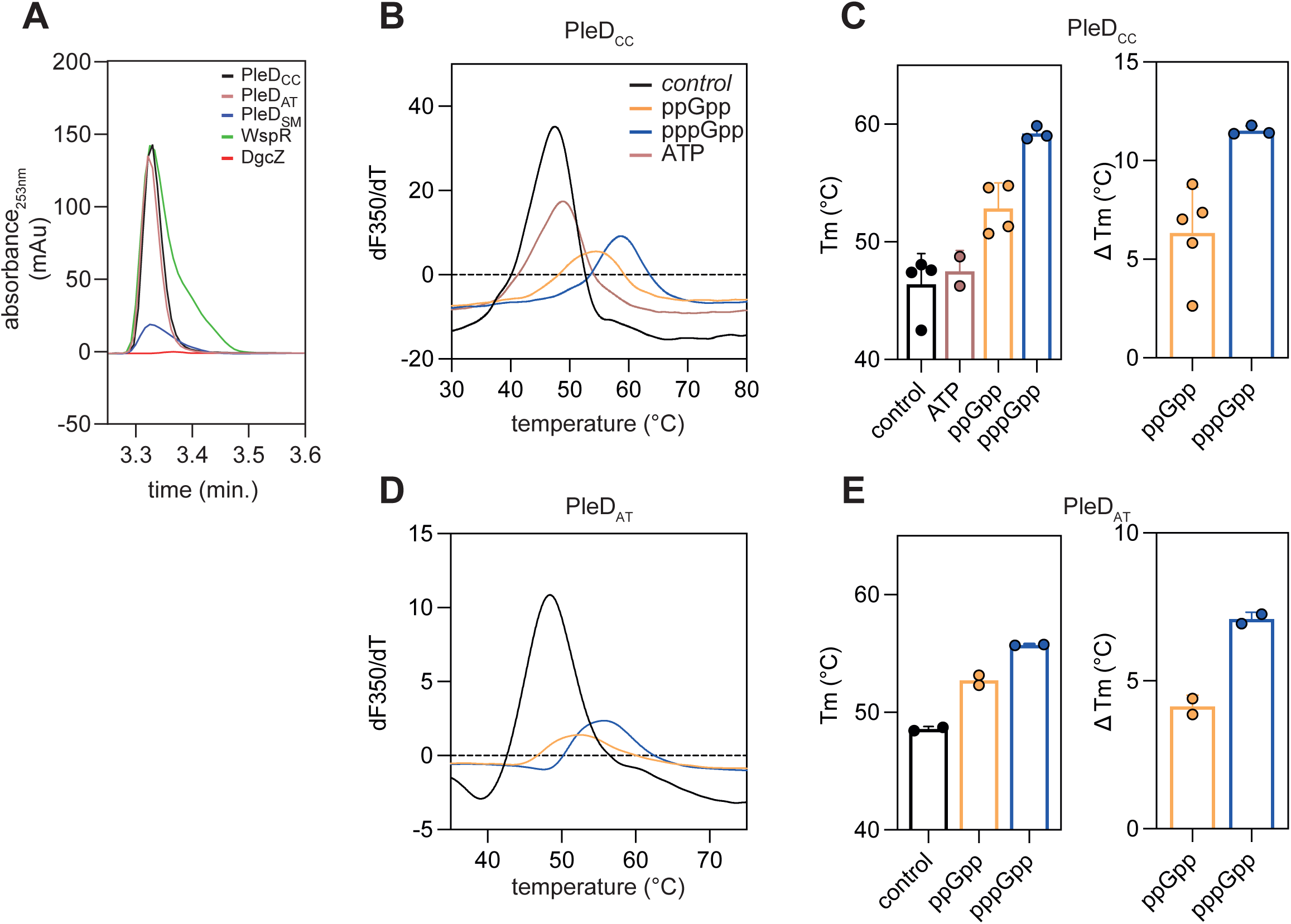
(A) HPLC profiles of different DGCs following purification from *E. coli* containing 40 µM PleD_CC_ (black), 40 µM PleD_AT_ (pink), 40 µM PleD_SM_ (blue), 40 µM WspR (green) or 40 µM DgcZ (red). (B) First-derivative curves at 350 nm from nanoDSF assays of 100 µM PleD from *Caulobacter crescentus* (PleD_CC_) without any ligand (black) or with 1 mM ATP (pink), 1 mM GDP (red), 1 mM ppGpp (orange), 1 mM GTP (green), or 1 mM pppGpp (blue). (C) Melting temperature (Tm) determination from nanoDSF assays of 100 µM PleD_CC_ (left). ΔTm values indicate the difference in Tm between PleD_CC_ without any ligand (black) and with GDP (red), ppGpp (orange), GTP (green), or pppGpp (blue). Data are presented as mean ± SD. (D) Same as (B) for 100 µM PleD from *Agrobacterium tumefaciens* (PleD_AT_). (E) Same as (C) for PleD_AT_.

**Figure S3 related to Figure 3:**
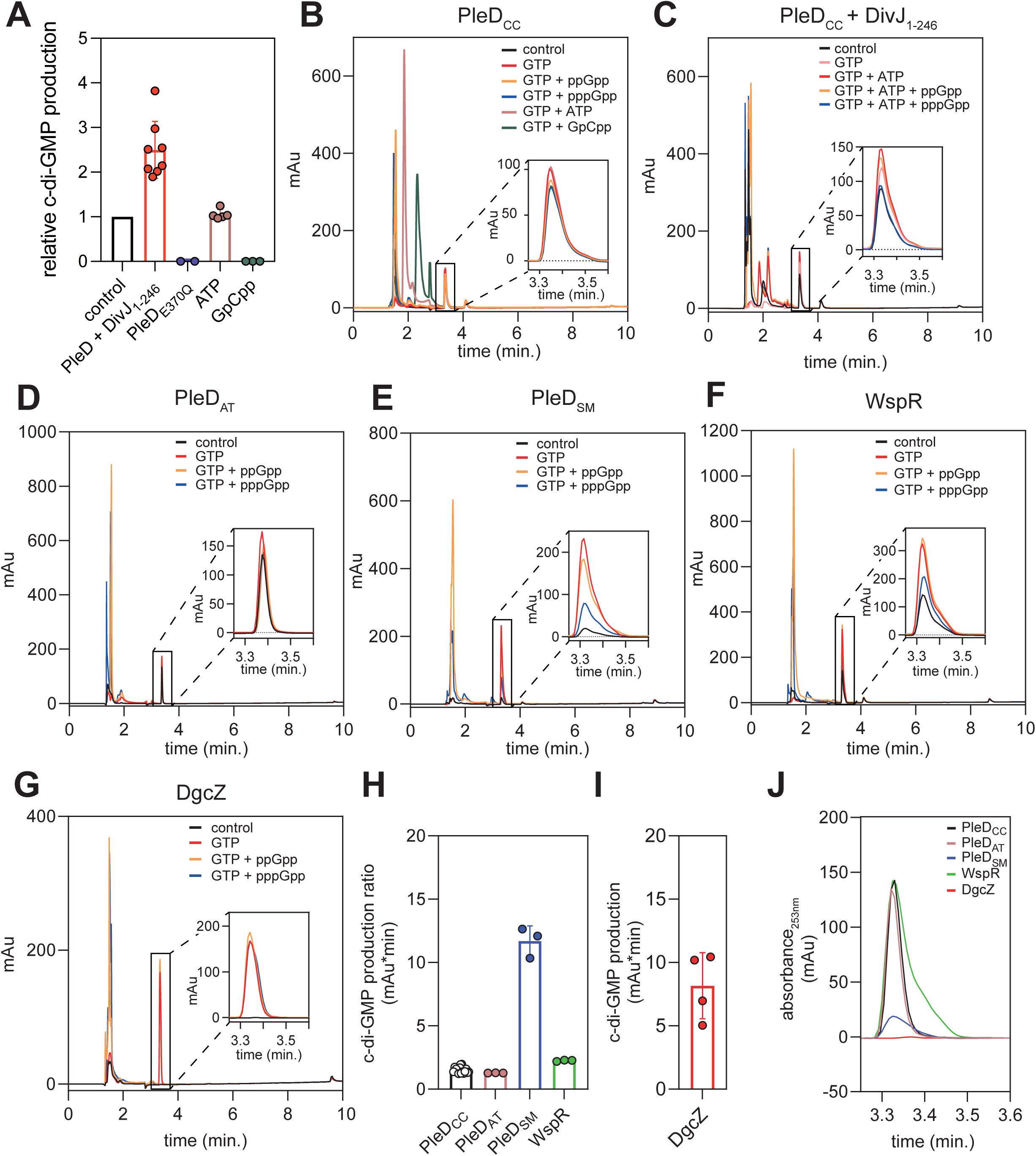
(A) *In vitro* enzymatic assay measuring c-di-GMP production by PleD or the catalytically-inactive PleD_E370Q_ mutant from *Caulobacter crescentus*. Reactions contain 40 µM PleD, 40 µM PleD pre-incubated with 10 µM DivJ_1-246_ and 200 µM ATP for 25 minutes or PleD_E370Q_. Bars represent 100 µM GTP alone (black), or with the addition of 500 µM ATP (pink), GpCpp (green), 100 µM GTP with PleD_E370Q_ (blue), or 100 µM GTP with phosphoactivated PleD (red). The relative c-di-GMP production represents the amount of c-di-GMP produced in each condition compared to the baseline production by PleD without competitor. Data are shown as mean ± SD. (B) High-performance liquid chromatography (HPLC) profiles of reaction mixtures containing 40 µM PleD alone (black), 40 µM PleD with 100 µM GTP (red), or 100 µM GTP with 500 µM ATP (pink), GpCpp (green), ppGpp (orange), or pppGpp (blue). A zoomed-in view of the specific c-di-GMP peaks (retention time ∼3.35 min) for each condition is highlighted in the black square. (C) Same as (B) for phospho-activated PleD pre-incubated with 10 µM DivJ_1-246_ and 200 µM ATP for 25 minutes. Reaction mixtures contain 40 µM PleD alone (black), 40 µM PleD with 100 µM GTP (pink), 40 µM phospho-activated PleD with 100 µM GTP (red), or 100 µM GTP with 500 µM ppGpp (orange), or pppGpp (blue). (D) Same as (B) for PleD ortholog from *Agrobacterium tumefaciens* (PleD_AT_). (E) Same as (B) for PleD ortholog from *Sinorhizobium meliloti* (PleD_SM_). (F) Same as (B) for WspR from *Pseudomonas aeruginosa*. (G) Same as (B) for DgcZ from *Escherichia coli*. (H) *In vitro* enzymatic assay measuring c-di-GMP production by different diguanylate cyclases (DGCs), presented as the ratio of c-di-GMP production with 100 µM GTP to the condition with protein only (not normalized). Data are presented as mean ± SD. (I) *In vitro* enzymatic assay measuring c-di-GMP production by DgcZ with 100 µM GTP. Data are presented as mean ± SD.

**Figure S4 related to Figure 4:**
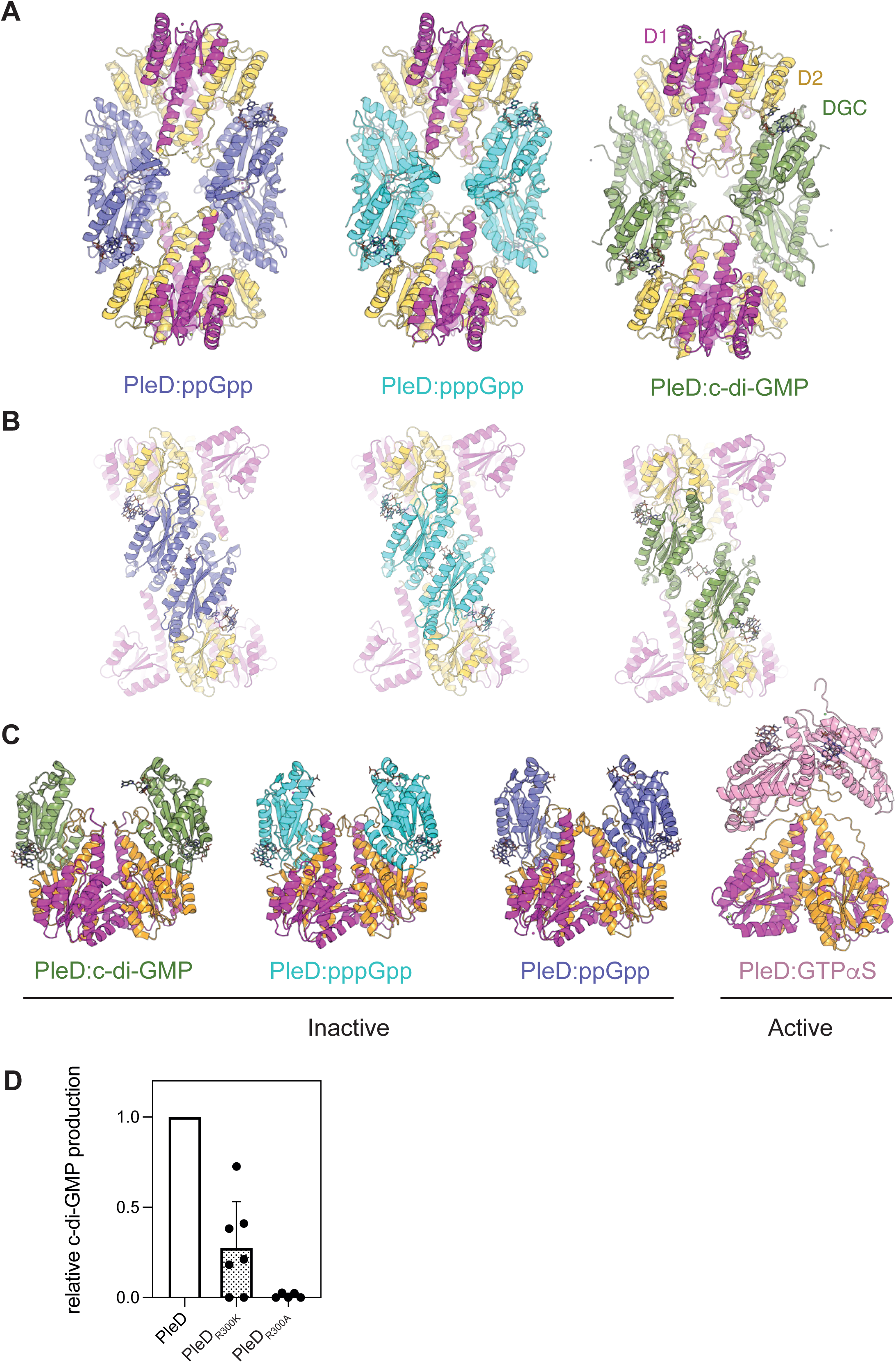
(A) Comparison of the PleD:c-di-GMP tetramer (PDB code 1W25, left) with the crystal structure of PleD in complex with ppGpp (middle) or pppGpp (right). Receiver domains D1 and D2 are colored in magenta and orange, DGC domains are colored according the crystal structure. (B) View of the dimer-dimer interface showing the two DGC domains facing each other. (C) Comparison of the dimer in « inactive » conformation and « activated » conformation in the different crystal structures colored as in (A). (D) *In vitro* enzymatic assay measuring c-di-GMP production by unphosphorylated PleD, PleD_R300K_, and PleD_R300A_. The reactions contained either 40 µM PleD (black), 40 µM PleD_R300K_ (dotted lines) or 40 µM PleD_R300A_ (dashed lines) with 100 µM GTP alone. Normalized c-di-GMP production is represented relative to the production by wild-type PleD.

**Figure S5 related to Figure 4:**
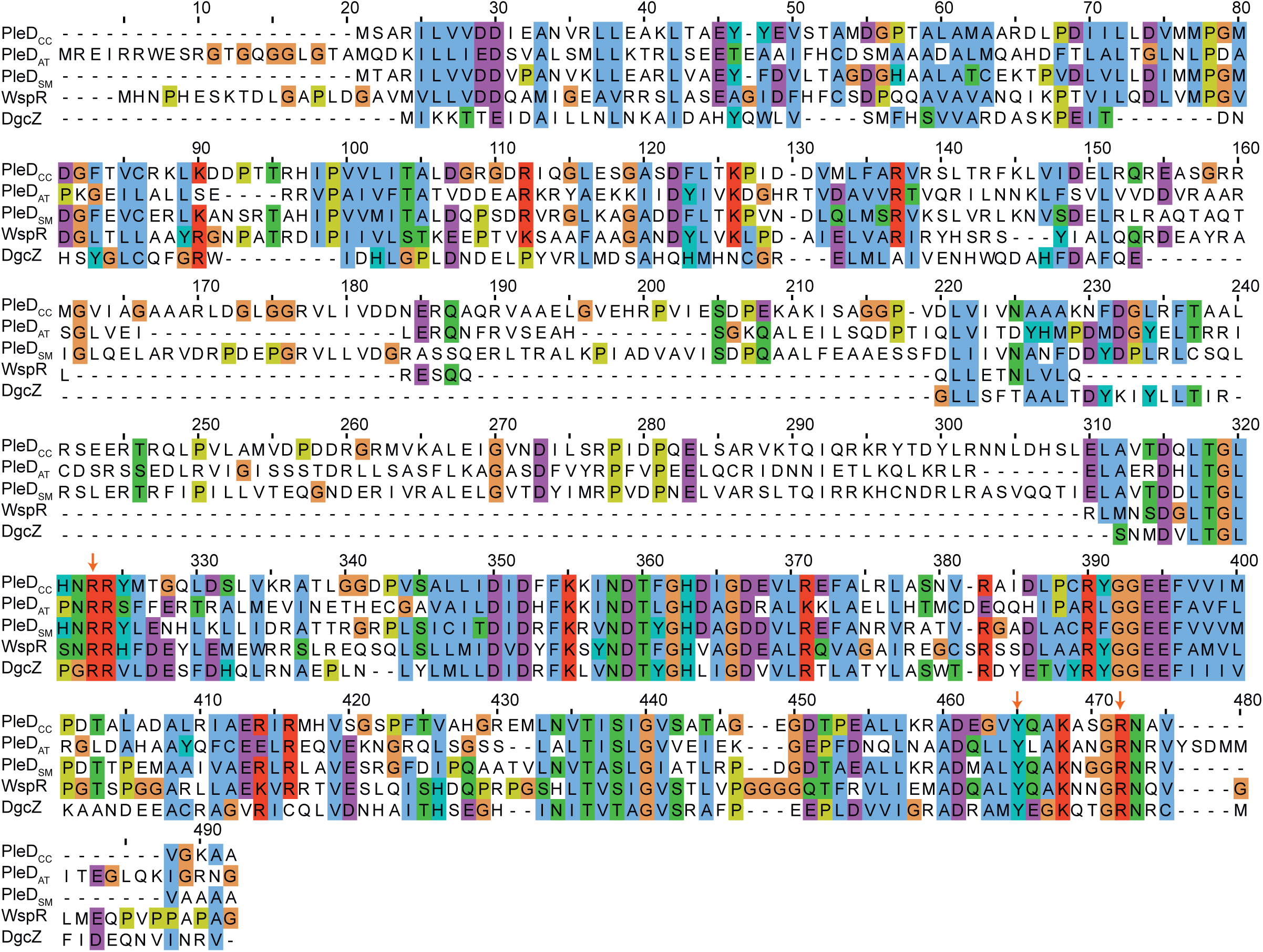
ClustalW sequence alignment of PleD_CC_, PleD_AT_, PleD_SM_, WspR and DgcZ. UNIPROT accession codes are B8GZM2, A9CJ98, Q92QM5, Q9HXT9 and P31129 respectively. Residue colors correspond to the Clustal color scheme: hydrophobic (blue), positively charged (red), negatively charged (magenta), polar (green), glycine (orange), proline (yellow), aromatic (cyan) and non-conserved (white) residues. Red arrows indicate highly conserved R300, Y439 and R446 residues.

**Figure S6 related to Figure 5:**
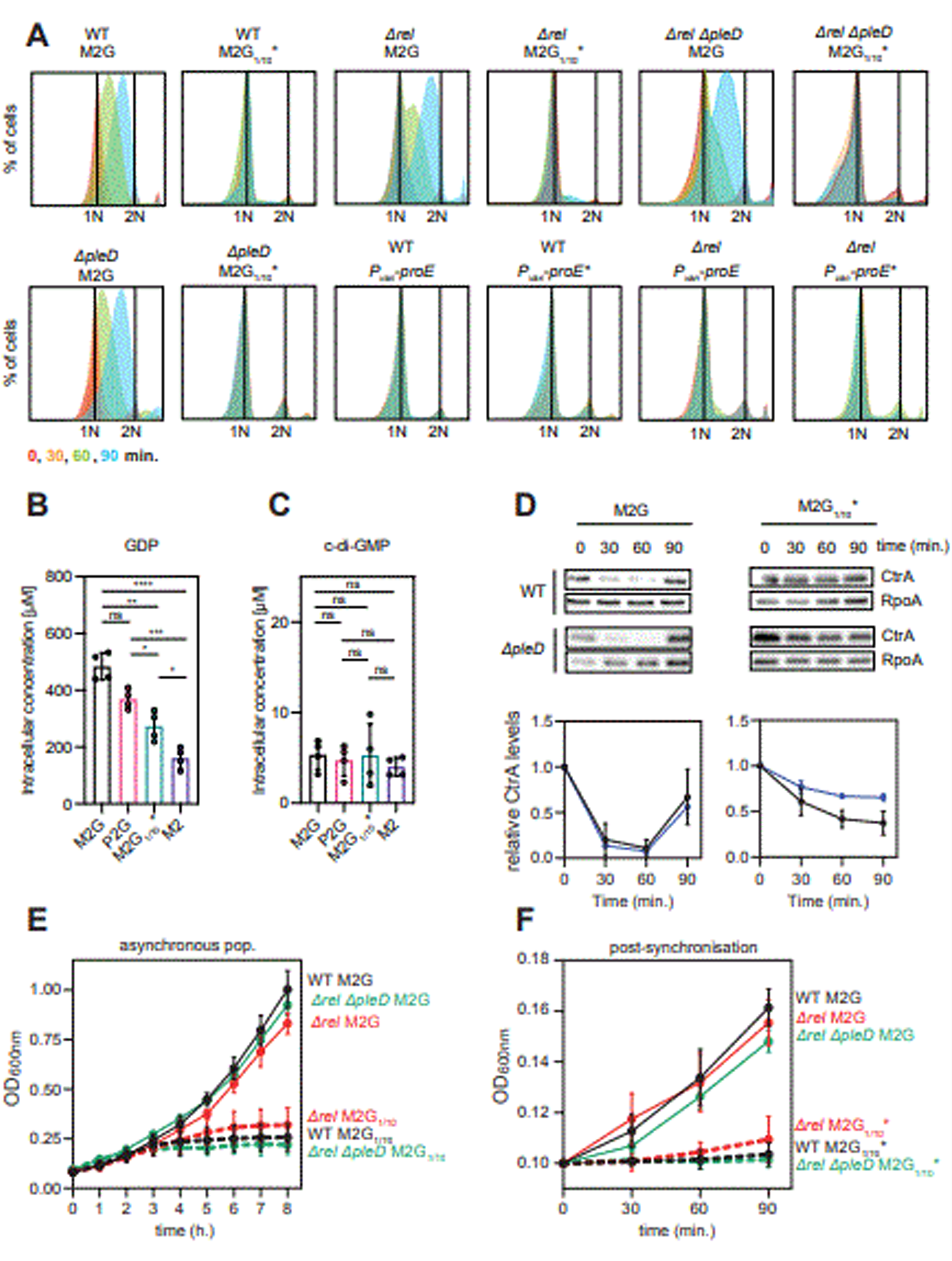
(A) Flow cytometry profiles after SYTOX staining showing DNA content of wild-type, *rel* mutant, *rel pleD* mutant and *pleD* mutant cells grown post-synchronization in M2 supplemented with 0.2% glucose (M2G) or 0.02% glucose spent medium (M2G_1/10_*) and of wild-type and *rel* mutant cells grown post-synchronization in M2 supplemented with 0.02% glucose spent medium with ectopic expression of *proE* or its inactive mutant *proE_E328A_* (induced with 100 µM vanillate post-synchronization). (B) Targeted metabolomics measuring the intracellular concentration of GDP in a asynchronous population of wild-type cells grown in M2G medium (black), P2G medium (pink), M2 with 0.02% glucose (M2G_1/10_*) (green) or M2 without glucose (purple). Data represent the mean ± SD of three independent biological replicates. Asterisks indicate statistical significance (** = p < 0.01; *** = p < 0.001) based on a Student’s t-test. (C) Same as (B) but representing c-di-GMP intracellular concentration. (D) CtrA immunoblots showing protein levels at the indicated times post-synchronization in wild-type and *pleD* mutant cells grown in supplemented with 0.2% glucose (M2G) or 0.02% glucose spent medium (M2G_1/10_*) over 90 minutes (left). Quantification of CtrA band intensity normalized to the RpoA loading control (right). Data represent mean ± SD of three independent biological replicates. Asterisks indicate statistical significance (* = p < 0.05; ** = p < 0.01) based on a Student’s t-test. (E) Growth curves of asynchronous wild-type, *rel* mutant, and *rel pleD* mutant cells cultivated in M2 supplemented with 0.2% glucose (M2G) or 0.02% glucose (M2G_1/10_*) over 8 hours. Data represent mean ± SD of three independent biological replicates. (F) Growth curves of wild-type, *rel* mutant, and *rel pleD* mutant cells cultivated post-synchronization in M2 supplemented with 0.2% glucose (M2G) or 0.02% glucose spent medium (M2G_1/10_*) over 90 minutes. Data represent mean ± SD of three independent biological replicates.

**Table S1:** Fitting results of ITC experiments.

| ligand | [ligand]<br>($\mu\text{M}$) | [PleD]<br>($\mu\text{M}$) | ratio | Chi<sup>2</sup> | N (sites) | K ($\text{M}^{-1}$) | Kd (M) | $\Delta\text{H}$<br>(cal/mol) | $\Delta\text{S}$<br>(cal/mol/deg) |
| --- | --- | --- | --- | --- | --- | --- | --- | --- | --- |
| GpCpp | 1000 | 95 | 10,5-1 | 1.06E+03 | 0.8711 | 2.22E+05 | 4.50E-06 | 1442 | 29.28 |
| GpCpp | 1000 | 80 | 12,5-1 | 725.44 | 0.9047 | 1.33E+05 | 7.53E-06 | 1273 | 27.71 |
| ppGpp | 800 | 100 | 8-1 | 2145 | 0.95 | 6.28E+04 | 1.59E-05 | 2993 | 31.99 |
| ppGpp | 1000 | 100 | 10-1 | 619 | 0.9 | 1.43E+04 | 6.99E-05 | 3586 | 31.04 |
| ppGpp | 1000 | 80 | 10-1 | 414 | 0.875 | 1.98E+04 | 5.05E-05 | 3008 | 29.75 |
| pppGpp | 800 | 80 | 10-1 | 2.52E+04 | 0.8769 | 9.76E+04 | 1.02E-05 | 3183 | 33.5 |
| pppGpp | 1000 | 100 | 10-1 | 1.37E+04 | 1.18 | 1.00E+05 | 1.00E-05 | 4383 | 37.6 |
| pppGpp | 750 | 75 | 10-1 | 3.12E+03 | 0.95 | 1.30E+05 | 7.67E-06 | 2506 | 31.81 |

**Table S2:** NanoDSF measurements.

| **T<sub>m</sub> (°C)** | **control** | **ppGpp** | **pppGpp** | **ATP** |
| --- | --- | --- | --- | --- |
| PleD *C. crescentus* | 46,39 ± 2,615 | 52,85 ± 2,146 | 59,21 ± 0,5707 | 47,5 ± 1,768 |
| PleD *A. tumefaciens* | 48,58 ± 0,2121 | 52,72 ± 0,601 | 55,72 ± 0,05657 |  |

| **Δ T<sub>m</sub> (°C)** | **ppGpp** | **pppGpp** |
| --- | --- | --- |
| PleD *C. crescentus* | 6,332 ± 2,323 | 11,52 ± 0,2318 |
| PleD *A. tumefaciens* | 4,135 ± 0,3815 | 7,092 ± 0,2275 |

**Table S3:**
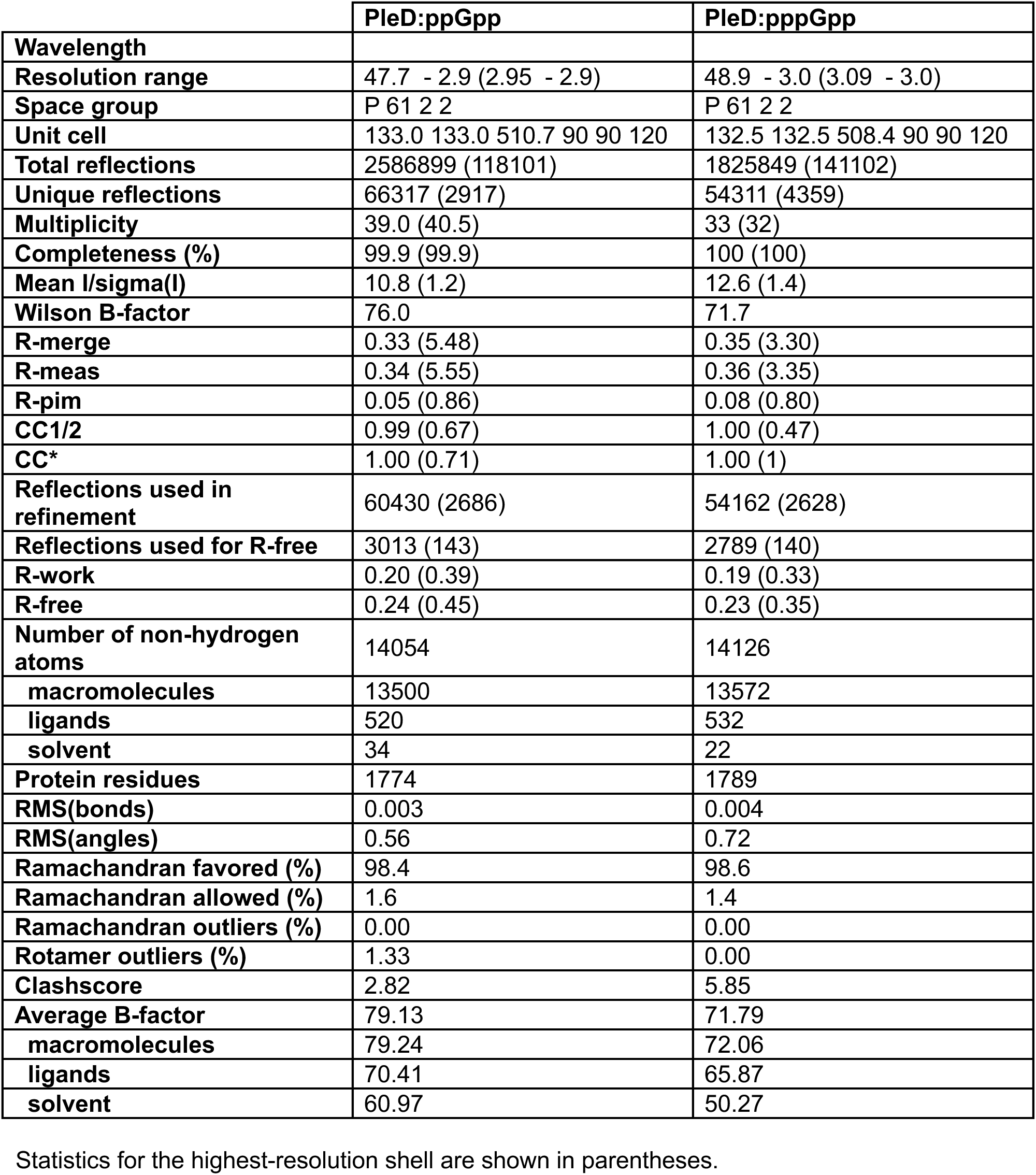
Data collection and refinement statistics.

|  | **PleD:ppGpp** | **PleD:pppGpp** |
| --- | --- | --- |
| **Wavelength** |  |  |
| **Resolution range** | 47.7 - 2.9 (2.95 - 2.9) | 48.9 - 3.0 (3.09 - 3.0) |
| **Space group** | P 61 2 2 | P 61 2 2 |
| **Unit cell** | 133.0 133.0 510.7 90 90 120 | 132.5 132.5 508.4 90 90 120 |
| **Total reflections** | 2586899 (118101) | 1825849 (141102) |
| **Unique reflections** | 66317 (2917) | 54311 (4359) |
| **Multiplicity** | 39.0 (40.5) | 33 (32) |
| **Completeness (%)** | 99.9 (99.9) | 100 (100) |
| **Mean I/sigma(I)** | 10.8 (1.2) | 12.6 (1.4) |
| **Wilson B-factor** | 76.0 | 71.7 |
| **R-merge** | 0.33 (5.48) | 0.35 (3.30) |
| **R-meas** | 0.34 (5.55) | 0.36 (3.35) |
| **R-pim** | 0.05 (0.86) | 0.08 (0.80) |
| **CC1/2** | 0.99 (0.67) | 1.00 (0.47) |
| **CC\*** | 1.00 (0.71) | 1.00 (1) |
| **Reflections used in refinement** | 60430 (2686) | 54162 (2628) |
| **Reflections used for R-free** | 3013 (143) | 2789 (140) |
| **R-work** | 0.20 (0.39) | 0.19 (0.33) |
| **R-free** | 0.24 (0.45) | 0.23 (0.35) |
| **Number of non-hydrogen atoms** | 14054 | 14126 |
| **macromolecules** | 13500 | 13572 |
| **ligands** | 520 | 532 |
| **solvent** | 34 | 22 |
| **Protein residues** | 1774 | 1789 |
| **RMS(bonds)** | 0.003 | 0.004 |
| **RMS(angles)** | 0.56 | 0.72 |
| **Ramachandran favored (%)** | 98.4 | 98.6 |
| **Ramachandran allowed (%)** | 1.6 | 1.4 |
| **Ramachandran outliers (%)** | 0.00 | 0.00 |
| **Rotamer outliers (%)** | 1.33 | 0.00 |
| **Clashscore** | 2.82 | 5.85 |
| **Average B-factor** | 79.13 | 71.79 |
| **macromolecules** | 79.24 | 72.06 |
| **ligands** | 70.41 | 65.87 |
| **solvent** | 60.97 | 50.27 |

Statistics for the highest-resolution shell are shown in parentheses.

**Table S4:** Calculated concentrations of GTP, GDP, ppGpp, pppGpp, c-di-GMP in *C. crescentus* grown in different media.

| Medium | GTP [$\mu\text{M}$] | GDP [$\mu\text{M}$] | ppGpp [$\mu\text{M}$] | pppGpp [$\mu\text{M}$] | c-di-GMP [$\mu\text{M}$] |
| --- | --- | --- | --- | --- | --- |
| M2G | 440,24 ± 50,447 | 483,76 ± 48,091 | 29,51 ± 8,815 | 4,60 ± 1,551 | 5,35 ± 1,610 |
| P2G | 378,46 ± 42,388 | 368,62 ± 34,331 | 85,51 ± 21,577 | 4,04 ± 0,937 | 4,72 ± 1,729 |
| M2G<sub>1/10</sub>* | 271,67 ± 30,445 | 273,62 ± 52,032 | 37,92 ± 8,035 | 10,37 ± 2,948 | 5,27 ± 3,449 |
| M2 | 119,12 ± 43,393 | 161,14 ± 36,616 | 55,43 ± 20,673 | 4,37 ± 1,799 | 4,02 ± 1,017 |

Data represent mean ± SD of four independent biological replicates.

**Table S5:**
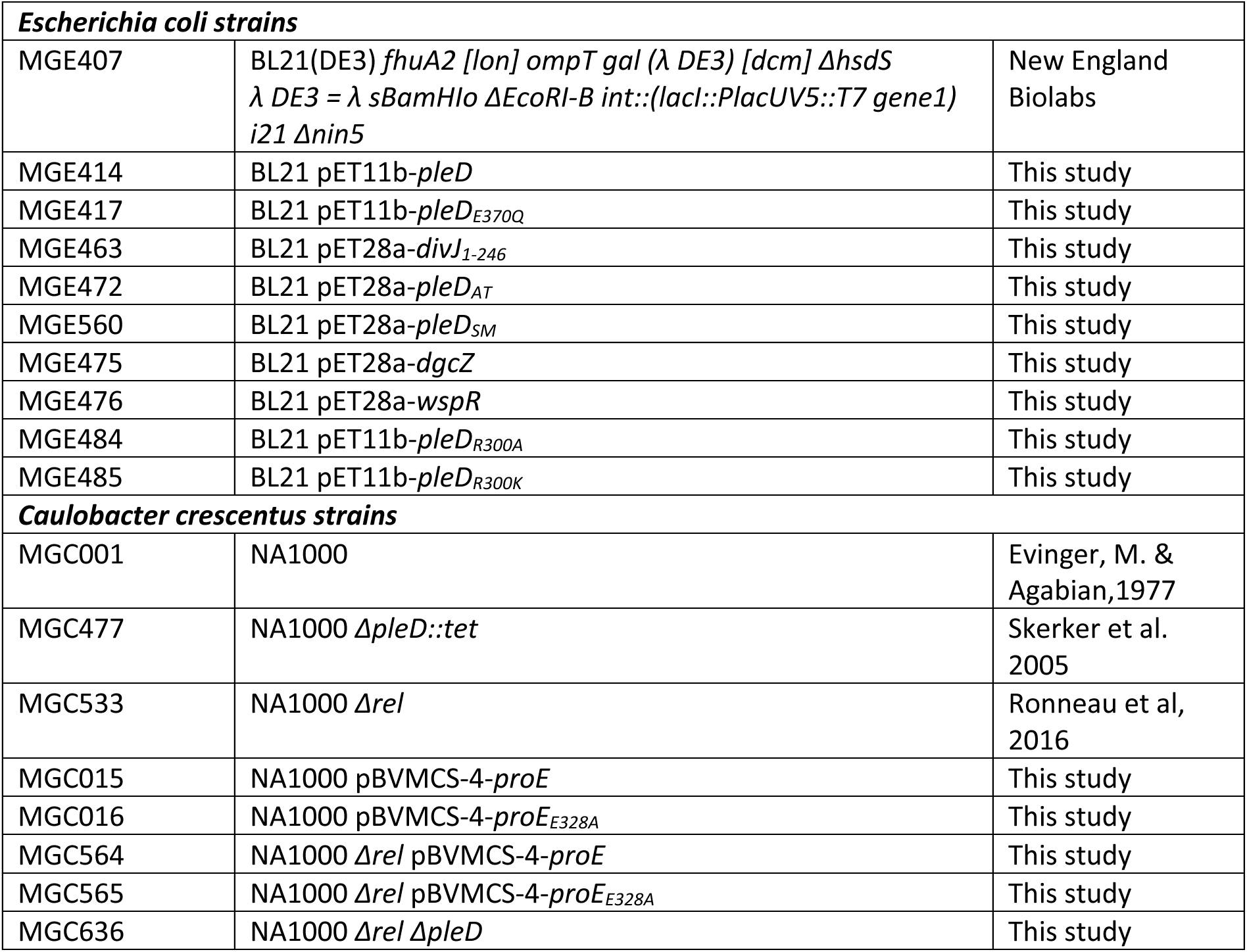
Bacterial strains used in this study.

**Table S6:** Plasmids used in this study.

|  |  |  |
| --- | --- | --- |
| MGE477 | pNTPS138-$\Delta spoT$ | Ronneau et al, 2016 |
| MGE351 | pBVMCS-6 | Thanbichler et al, 2007 |
| MGE003 | pBVMCS-4-*proE* | Duerig et al, 2009 |
| MGE004 | pBVMCS-4-*proE*<sub>E328A</sub> | Duerig et al, 2009 |
| MGE409 | pET11b-*pleD* | Paul et al, 2004 |
| MGE412 | pET11b-*pleD*<sub>E370Q</sub> | Paul et al, 2007 |
| MGE406 | pET28a | Novagen |
| MGE506 | pNTPS138-*pleD* | This study |
| MGE462 | pET28a-*divJ*<sub>1-246</sub> | This study |
| MGE470 | pET28a-*pleD*<sub>AT</sub> | This study |
| MGE473 | pET28a-*dgcZ* | This study |
| MGE474 | pET28a-*wspR* | This study |
| MGE481 | pET11b-*pleD*<sub>R300A</sub> | This study |
| MGE482 | pET11b-*pleD*<sub>R300K</sub> | This study |
| MGE643 | pBVMCS-6-*dgcZ* | This study |

**Table S7:**
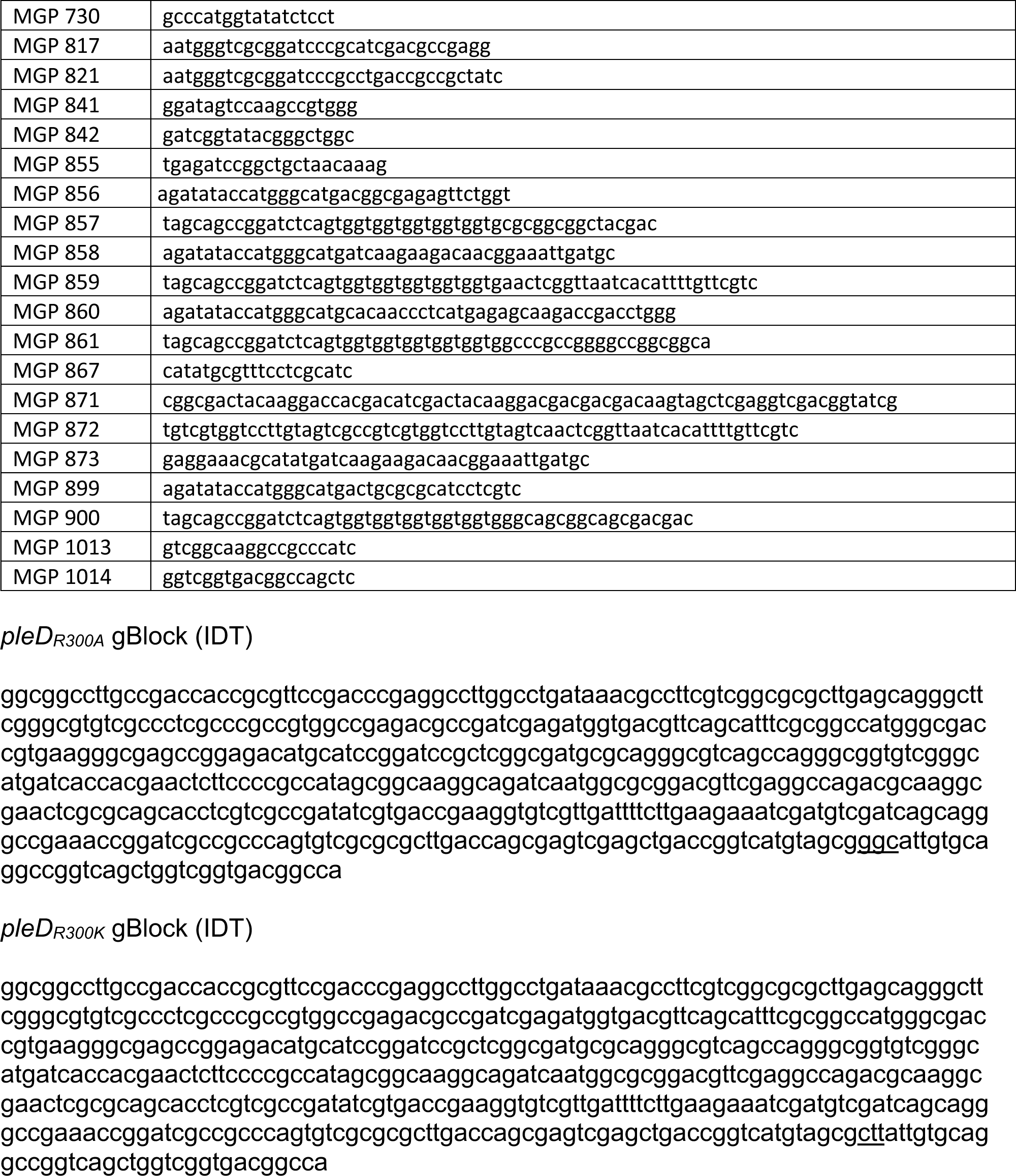
Oligonucleotides and synthetic DNA sequences used in this study.

## Notes

### Competing Interest Statement

The authors have declared no competing interest.

